# Artificial Reactivation of a Cocaine-Associated Ensemble in the Dorsal Dentate Gyrus Attenuates Cocaine Prime-Induced Reinstatement of Conditioned Place Preference

**DOI:** 10.64898/2026.05.19.726387

**Authors:** LH Edwards, LF Papanikolaou, MR Wilson, MV Brody, WF Wade, M Culter, SA Arora, A Stratmann, S Cañuelas del Valle, SL Grella

## Abstract

Relapse-prevention strategies aimed at reducing relapse following abstinence primarily focus on reducing cravings that lead to drug-seeking triggered by stress, drug-related cues, or re-exposure to the drug. Because addictive drugs form persistent associative contextual memories, we investigated how reactivation of cocaine-related hippocampal memories influences subsequent drug-seeking. Here, we tagged dorsal dentate gyrus (dDG) memory ensembles involved in encoding either a first or fourth cocaine exposure (15mg/kg, i.p) in male and female c57BL/6 mice using a TetTag approach. Mice underwent cocaine conditioned place preference (CPP), extinction, and reinstatement. We assessed whether optical reactivation of tagged cocaine-related ensembles could substitute for a cocaine priming injection to reinstate CPP, whether reactivation altered cocaine-induced reinstatement, and if these effects differed depending on stage of drug exposure. We also compared these effects to reactivation of saline-associated ensembles. Cocaine produced robust locomotor activation during conditioning and sensitization developed across repeated drug exposures. Reactivation of a cocaine-related engram alone did not reinstate CPP. However, reactivation of the first cocaine exposure engram attenuated cocaine-induced reinstatement. In contrast, reactivation of the fourth exposure engram did not confer this protective effect. Interestingly, reactivation of saline-associated ensembles also reduced cocaine-induced reinstatement specifically in females, suggesting dDG ensemble reactivation may modulate relapse-related behavior through interference or neuromodulatory disruption of cocaine-associated representations, consistent with our prior work. These findings raise the possibility that early contextual experiences form competing or destabilizing representations that interfere with later cocaine-seeking when reactivated. Females also displayed greater sensitivity to locomotor-inducing effects of cocaine memory reactivation, although this was dissociated from CPP. Together, these findings show that cocaine memories are distinct across drug experience and selective reactivation of dDG engrams can differentially influence drug-seeking.

**Graphical Abstract Text:** Activity-dependent tagging was used to label dorsal dentate gyrus (dDG) neuronal ensembles recruited during either the first or fourth cocaine exposure in a cocaine conditioned place preference (CPP) task. Following training and extinction, tagged ensembles were optogenetically reactivated during cocaine-primed reinstatement. Reactivation of neurons tagged during the first cocaine exposure attenuated reinstatement, whereas this effect was not observed in animals tagged during the fourth exposure. These findings suggest that neuronal ensembles recruited during early cocaine experiences can influence subsequent cocaine-seeking behavior.

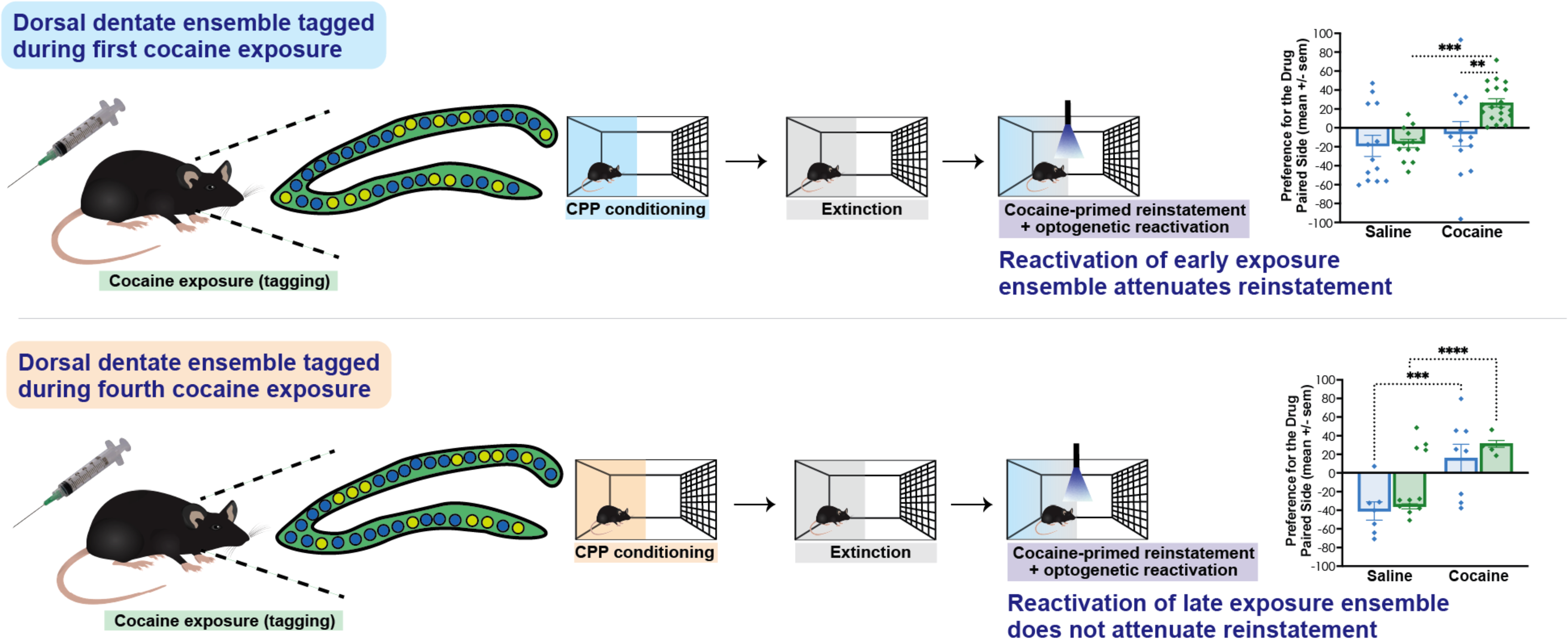

## 1. INTRODUCTION

Substance Use Disorders represent a major public health challenge with relapse remaining one of the greatest obstacles to treat (Larimer et al., 1999; Nestler, 2025). Drug-associated cues, contexts, stress, and re-exposure to the drug can all trigger drug-seeking behavior following abstinence (Martin-Fardon & Weiss, 2013; Shaham et al., 2003). Accordingly, relapse-prevention research has focused heavily on the neural circuits underlying cue-, stress-, and drug-induced reinstatement using the reinstatement model (de Wit & Stewart, 1981, 1983; Koob et al., 2009; Lintas et al., 2011; Mantsch et al., 2016; McFarland & Kalivas, 2001). While these studies have identified important roles for the amygdala, nucleus accumbens, prefrontal cortex, and ventral tegmental area, the contribution of drug-related memories and hippocampal circuits to reinstatement remains less understood. Given the established role of the dorsal dentate gyrus (dDG) in contextual memory (Cutler et al., 2025; Grella et al., 2016, 2022; Grella & Donaldson, 2024; Maple et al., 2017; Smith & Bulkin, 2014), we were interested in whether artificial reactivation of a drug-related memory recapitulates the drug experience in a manner similar to the drug (i.e., promoting relapse like a priming injection) or does it function more like extinction (i.e., attenuating relapse) where a conditioned stimulus is presented in the absence of the unconditioned stimulus? Addressing these questions may provide insight into how contextual drug memories contribute to cocaine-seeking behavior and relapse vulnerability (Cutler et al., 2025).

The use of viral neuronal labeling strategies such as the Tet Tag system makes it possible to genetically label and then optically reactivate memories using optogenetics (Grella et al., 2022; Reijmers et al., 2007). This system is both activity-dependent and inducible. Its activity dependence stems from its reliance on the immediate early gene c-Fos as a promoter, while its inducibility allows precise temporal control over tagging windows using doxycycline (DOX, 40 mg/kg food pellets). When mice are maintained on DOX, the system remains inactive. However, replacing DOX with regular chow opens a tagging window, triggering the expression of channelrhodopsin-2 (ChR2) fused to a fluorescent reporter (e.g., enhanced yellow fluorescent protein, eYFP). This enables both targeted manipulation of tagged dDG cells and their visualization under a microscope.

Using this approach, in experiment 1, we trained C57BL/6 mice on cocaine conditioned place preference (CPP), a widely used and effective model for examining the association between contextual cues and drug-related experiences (McKendrick & Graziane, 2020) and tagged dDG encoding their first exposure to cocaine on the initial day of conditioning. In this procedure, mice learn to associate one side of the apparatus with cocaine and the other with saline, resulting in a preference for the cocaine-paired side due to the rewarding effects of the drug. After this preference is extinguished, it can be reinstated by a cocaine priming injection (Mueller & Stewart, 2000). Here, we tagged hippocampal ensembles active during the initial cocaine exposure on the first day of conditioning and later tested whether reinstatement following extinction could be induced by either a cocaine prime or optical reactivation of the cocaine-associated memory as well as both. Our previous work demonstrated that tagging and reactivating dDG cells involved in encoding an acute first exposure to cocaine did not produce locomotor effects comparable to those induced by the drug (Grella et al., 2022). While unsurprising, given that the dDG is not typically linked to locomotion, this finding raises the question: what aspects of the drug experience are being recapitulated? To explore this further, in experiment 2, we compared locomotor responses in an open field following reactivation of engrams tagged during either the first or fourth cocaine exposure. Then, in experiment 3, we returned to the original experimental design used in experiment 1 but shifted the tagging window such that dDG neurons active during the fourth cocaine exposure, rather than the first, were labeled and subsequently reactivated optically. Finally, in experiment 4, we tagged a saline exposure on the first day of conditioning to control for nonspecific effects of neuronal tagging and optical reactivation independent of drug experience. Leveraging viral neuronal tagging strategies in this way, we hope to gain insights into the mechanisms underlying addiction-related pathways, which may pave the way for novel relapse-prevention strategies.

## 2. EXPERIMENTAL PROCEDURES

### 2.1 Animals

All experimental procedures were approved by the Institutional Animal Care and Use Committee at Loyola University Chicago and conducted in accordance with the NIH Guide for the Care and Use of Laboratory Animals. The experimental mice, consisting of c57BL/6 mice (∼39 days old at the experiment’s onset; Charles River Labs, 027) had an average weight of 21-25g for males and 18–22g for females upon arrival. Mice were housed in groups of 2-3 per cage in a temperature and humidity-controlled colony room (temperature: 18-25°C, humidity: 40-70%), following a regular 12:12 h light ON/OFF cycle. Experiments were conducted during the light phase of the cycle. Cages, equipped with huts and nesting material for enrichment, were changed weekly. Upon arrival at the facility, all mice were placed on a 40 mg/kg DOX diet (Bio-Serv, product F4159, Lot 28189) and were left undisturbed for a minimum of 3 days before surgery, with *ad libitum* access to food and water. Additionally, mice were handled for two consecutive days for 2 minutes each, and on the third day, they were both handled and received a saline injection to acclimate them to the injection procedure.

### 2.2 Stereotaxic surgery

Aseptic surgeries were conducted in a stereotaxic frame (Kopf Instruments) with the skull positioned flat and mice resting on a heating pad. They were initially anesthetized with 4% isoflurane and 80% oxygen (induction), followed by a reduction to 2% isoflurane (maintenance). At the start of surgery, mice received 0.2 mL physiological sterile saline (0.9%, s.c.) and carprofen (10 mg/kg, i.p.) and ophthalmic ointment over their eyes. The surgical site was prepped by removing the hair with Nair, and then swabbing with 95% isopropyl alcohol solution, betadine, and lidocaine with 2% epinephrine. An incision was made, and the skin was pulled back with two bulldog clamps. Viral infusions were delivered using a 10 μL gas-tight Hamilton syringe connected to a micro-infusion pump (UMP3, World Precision Instruments) at a rate of 100 nL/min. The infusion needle remained in place for 10 minutes post-infusion to prevent liquid backflow. Optic fibers were implanted, and secured with two anchor screws, and a head cap was constructed using a mixture of metabond (Parkell) and dental cement (Kopf). Post-surgery, mice were placed on a heating pad, provided with hydrogel, and given *ad libitum* access to food and water. An additional injection of carprofen (10 mg/kg i.p.) was administered the next day, followed by an injection of meloxicam (5 mg/kg, s.c.) 24 hours later. Apart from cage changes and health monitoring, mice were left undisturbed for a 7–10-day period post-surgery to facilitate recovery and allow for virus expression.

### 2.3 Viral microinjections

To label cells involved in a cocaine or saline experience, mice underwent bilateral infusions of a viral cocktail, which consisted of two vectors: pAAV9-cFos-tTa (UMass Vector Core - titer: 2.7x10^13 GC/mL) and pAAV9-TRE-(ChR2)-eYFP (UMass Vector Core - ChR2 & eYFP titer: 1.6x10^13 GC/mL). Each infusion was delivered at a volume of 400 nL per side. The infusions were targeted at the dDG (AP: -2.2, ML: ±1.3, DV: −2.0 relative to Bregma in mm). Additionally, bilateral optic fibers were implanted at coordinates AP: -2.2, ML: ±1.3, DV: −1.6 (relative to Bregma in mm).

### 2.4 Genetically labelling dDG neurons involved in a behavioral epoch

We employed an activity-dependent and inducible Tet-Tag (tetracycline-inducible) optogenetic system to genetically label neurons that were active during distinct behavioral epochs (Doucette et al., 2020; Grella et al., 2022; Reijmers et al., 2007). This system utilizes the delivery of an adeno-associated viral (AAV) cocktail, enabling the expression of a tetracycline transactivator protein (tTA) and a tetracycline response element (TRE). When bound, the latter facilitated the expression of the light-sensitive protein channelrhodopsin-2 (ChR2), fused to the fluorescent reporter gene, enhanced yellow fluorescent protein (eYFP). The system was inducible because transcription could be reversibly turned on or off by tetracycline or its more stable derivative, doxycycline (DOX), which was present in the animal’s diet. To permit activity-dependent labeling, DOX-containing chow was replaced with standard chow (*ad libitum*) 42 h prior to the targeted behavioral session. This approach allowed for the inducible and activity-dependent expression driven by the c-Fos promoter, widely recognized as a neuronal marker (Chung, 2015). Specifically, we labeled the cells in the dDG that were active during an acute cocaine or saline-associated experience. After behavioral tagging, mice were returned to their home cages and once again placed on the DOX diet.

Importantly, removal of DOX chow does not result in an immediate onset of activity-dependent labeling, as residual DOX must decline sufficiently to permit tTA-dependent transcription from the TRE (Reijmers et al., 2007). Thus, the 42-h off-DOX interval includes the time required for DOX clearance and does not represent 42 h of equivalent labeling. Animals remained undisturbed in their home cages during this interval until the experimental session. Although labeling was targeted to the behavioral session, activity associated with experimental procedures on the day of tagging, including transport, handling, and injection, may also contribute to the labeled population.

### 2.5 Tagged acute cocaine (or saline) exposure

Mice were habituated to intraperitoneal (i.p.) saline injections on the final day of handling. Cocaine hydrochloride was dissolved in 0.9% NaCl to a concentration of 2.5 mg/ml, supplied by the NIH NIDA Intramural Research Drug Supply Program. On the day of behavioral tagging, mice received an injection of either cocaine (15 mg/kg, i.p.) or saline (0.15 ml).

### 2.6 Reactivating dDG neurons involved in a behavioral epoch

Optical stimulation of the genetically labeled dDG ensemble was delivered at 20 Hz with a 10-ms pulse width. Optic fiber implants were connected to patch cords coupled to a blue laser diode (473 nm; Doric Lenses) controlled using Doric Neuroscience Studio (version 6.2.4.0). Laser power was measured with an energy meter (Thorlabs) at the outset of each experiment to ensure a minimum output of 10 mW at the end of the patch cord. In addition, each optic fiber was tested prior to surgery to verify a minimum power output of 5 mW at the fiber tip.

Mice were connected to the patch cords immediately before placement into the CPP apparatus. Optical stimulation began upon placement into the apparatus, simultaneously with the start of the CPP test, and continued throughout the 10-min session. During cocaine-primed reinstatement testing, mice received cocaine (7.5 mg/kg, i.p.) immediately before placement into the CPP apparatus; optical stimulation therefore began immediately following cocaine administration and continued throughout the 10-min reinstatement test. (Lau et al., 2020)

### 2.7 Experimental design and procedures

#### 2.7.1 CPP (experiment 1): first cocaine exposure tagged

The behavioral assessments were conducted within conditioning chambers (dimensions: 59.7L x 30W x 34H cm) equipped with centrally positioned web cameras. Each chamber was divided into three distinct sections labeled as "dark," "middle," and "light." The dark compartments featured various color options (blue, black, or green) with patterned walls and a dark gray patterned floor. In contrast, the light sides were characterized by white walls without patterns and white textured floors. The middle regions served as transition areas with one wall light and the other dark and the flooring smooth and light gray. After each session, subjects were placed in a holding cage and not returned to their home cage until all cage mates underwent testing, upon which they were reunited. Subsequently, locomotor and behavior analyses were conducted using Ethovision XT 17 software. The CPP procedure spanned 10 days, with each session lasting 10 minutes.

#### 2.7.2 Dependent variables

For all sessions, locomotor measures were taken - mean speed (m/s) and distance traveled (m) were recorded. For test sessions, time spent in each compartment (light, middle and dark) was measured and then used to calculate preference ratio (%), preference score, and the CPP change score. Preference Ratio: Time on Side A / (Time on Side A - Time on Side B) *100. Preference Score: % of Time on Side A - % of Time of Side B. CPP Change Score: % of Time on Side A (Test2) - % of Time on Side A (Test 1).

#### 2.7.3 Pretests

To assess whether mice had an initial preference for one compartment over another, mice were given 3 pretest sessions (10 minutes each) where they were first placed in the middle compartment for 2 min and followed by access to the entire chamber for 10 minutes. Data from these 3 tests were averaged. We used a biased design, pairing the cocaine injections with placement in the light side as all mice preferred the dark side.

#### 2.7.4 Conditioning

On conditioning days 2, 4, 6, 8, and 10, mice were administered saline injections (0.15ml, i.p.) just before the commencement of each session. Conversely, on conditioning days 1, 3, 5, 7, and 9, mice received cocaine injections (15mg/kg, i.p.) immediately before each session. Mice were confined to only one side of the chamber throughout conditioning (dark = saline; light = cocaine).

#### 2.7.5 Conditioning test

Following 10 days of conditioning, mice were given a test to assess preference. They were placed in the center compartment for 2 minutes and then had access to the entire chamber for the duration of the 10-minute test.

#### 2.7.6. Extinction training

Mice then received 6 training sessions (10-minute sessions, 1 session per day). Mice were again confined to only one side of the chamber throughout extinction. However, they were given saline in both compartments (dark = saline; light = saline).

#### 2.7.7 Extinction test

Following 6 days of extinction, mice were given a test to assess preference. They were placed in the center compartment for 2 minutes and then had access to the entire chamber for the duration of the 10-minute test.

#### 2.7.8 Reinstatement test

The next day, mice were given a reinstatement test to assess preference. Prior to the test, they were either given an injection of saline (0.15 ml, i.p.) or cocaine (7.5 mg/kg, i.p.). All mice (eYFP and ChR2) were given light stimulation to reactivate the tagged cocaine-associated ensemble.

#### 2.7.9 Locomotor test (experiment 2)

We have previously shown that tagging dDG-mediated cells involved in the first exposure to cocaine (15mg/kg, i.p., - 45 minutes in a clean cage) does not induce locomotor effects when acutely reactivated (Grella et al., 2022) in a different context (in an open field). Because locomotor sensitization is highly context-specific (Mattson et al., 2008), we sought to test this more systematically by assessing whether artificial reactivation of a cocaine-associated memory can induce locomotion in the same context it was encoded in. To this end, we assigned mice to one of three open fields that differed in both wall patterns (striped, polka dot, or checkered) and floor textures (smooth rubber, ribbed rubber, or rubber with raised dots) and habituated them to this chamber for 5 minutes without drug on board to get a baseline measure of locomotion. Each open field measured 24” L x 12” W. Forty-eight hours later, we began giving mice cocaine injections (15mg/kg, i.p.), every other day across 8 days (4 total injections) followed immediately by placement into the open field for 30 minutes. Off-DOX, we tagged dDG neurons active during either the first or the fourth cocaine exposure and assessed locomotor behavior during acute memory reactivation (10 min; 20 Hz; 473 nm; 10 ms pulse width) in a 10-minute test session. Mice either received ChR2, eYFP, or no virus (mice received same surgery and optical implantation however no virus was injected into the brain) in the dDG and only the No Virus group received cocaine (7.5mg/kg, i.p.) immediately prior to the reinstatement test as a cocaine-induced sensitization positive control. Locomotion was quantified as mean speed recorded by analyzing videos through Ethovision XT.17.

#### 2.7.10 CPP (experiment 3): fourth cocaine exposure tagged

In Experiment 1, dDG neurons were tagged during the first cocaine exposure (Day 1). In Experiment 3, the experimental design was otherwise identical, but tagging occurred during the fourth cocaine exposure (Day 7).

#### 2.7.11 CPP (experiment 4): saline exposure tagged

In Experiment 4 (saline control), the conditioning schedule was reversed such that saline injections were administered on Days 1, 3, 5, 7, and 9, with the first saline exposure (Day 1) tagged. Cocaine injections (15 mg/kg, i.p.) were administered immediately before each session on Days 2, 4, 6, 8, and 10.

#### 2.7.12 Validation of DOX-dependent neuronal tagging (experiment 5)

To assess background labeling under DOX conditions, an additional cohort of mice (n = 5) was maintained on DOX throughout the experimental procedure. Brain tissue was collected and processed for eYFP and c-Fos visualization as described above. eYFP expression was examined in the dDG to assess background activity-dependent labeling in the presence of DOX.

#### 2.7.13 Histological validation of optogenetic reactivation (experiment 6)

To assess ChR2-dependent neuronal recruitment following optical stimulation, an additional cohort of mice expressing either ChR2-eYFP (n=5) or eYFP alone (n=5) underwent activity-dependent neuronal tagging during cocaine exposure. Mice were subsequently placed in a novel context and received identical optical stimulation (473 nm, 20 Hz, 10-ms pulses). Animals were perfused 90 min following stimulation, and brain sections were processed for c-Fos immunohistochemistry. The percentage of eYFP-positive neurons coexpressing c-Fos was quantified to assess recruitment of the tagged neuronal population following stimulation.

#### 2.7.14 Histology

Mice were transcardially perfused with 1x phosphate-buffered saline (PBS) at 4°C, followed by 4% paraformaldehyde solution (PFA). Brains were extracted and stored overnight in PFA at 4°C. They were subsequently transferred to a solution containing 0.01% sodium azide in 1x PBS the next day. Brains were sectioned into 50 μm coronal slices using a vibratome (Leica, VT100S) and collected in cold 1x PBS and placed in a well plate containing 0.01% sodium azide in 1x PBS and kept at 4°C.

#### 2.7.15 Immunohistochemistry

Sections were washed 3 times in 1x PBS (10 minutes each) and then blocked for 2 hours at room temperature in blocking solution (1x PBS + 2% Triton (PBS-T) and 5% normal goat serum) (NGS, Vector Laboratories) on an orbital shaker. This was followed by 2 days of incubation in blocking solution containing the primary antibody [1:1000 chicken anti-GFP (Invitrogen a10262)] kept on a shaker at 4°C. Afterward, the sections underwent 3 washes (10 minutes each) in PBS-T followed by a 2-hour incubation period at room temperature with blocking solution containing the secondary antibody prepared in PBS-T [1:200 Alexa 488 anti-chicken (Invitrogen, A11039)]. Following 3 more 10-minute washes in PBS-T, the sections were mounted onto microscope slides and counterstain was applied before coverslipping (DAPI added to Vectashield HardSet mounting medium, Vector Laboratories). Slides were placed at 4°C overnight to cure. The following day, the edges were sealed with clear nail polish, and the slides were stored in a slide box in the fridge until imaging.

#### 2.7.16. Fluorescent confocal image acquisition and quantification

Images were captured from coronal sections using a fluorescent confocal microscope (Zeiss LSM 880) at 20x magnification, and Zen (Black Edition) software was employed for acquisition. For the quantification of cells comprising cocaine-associated ensemble and to verify bilateral viral dDG injections, three z-stacks (step size 1 μm) were captured per hemisphere from three different slices, resulting in six z-stacks per animal. Data from each hemisphere were combined, and the means for the six z-stacks were calculated. These means were then used to derive a group mean. For all images, the total number of DAPI-positive (+) and eYFP+ neurons was quantified using Image J/Fiji software (https://imagej.nih.gov/ij/). This was done to verify injections but also to quantify the size of the cocaine-associated ensemble across groups.

#### 2.7.17 Statistical analysis

Experimenters were blinded to experimental group assignments during behavioral testing, histological quantification, and data analysis. Group identities were revealed only after data collection and scoring / counting were completed. Calculated statistics are presented as means ± standard error of the mean (s.e.m.). To assess differences, one, two, and three-way analysis of variance (ANOVA) was employed, and for cases involving repeated measures (RM) analyses, this is explicitly stated. Unpaired t-tests were utilized in some instances. Follow-up post-hoc comparisons (Tukey’s HSD) were conducted when deemed appropriate. All statistical analyses were performed using GraphPad Prism (version 9.2.0), were two-tailed, and assumed an alpha level of 0.05. In all figures, statistical significance is denoted as follows: *=p<0.05, **=p<0.01, ***=p<0.001, ****=p<0.0001. The individual animal served as the experimental unit for all behavioral and histological analyses. Mistargeted animals were excluded from the analysis, which resulted in 4 animals excluded in experiment 1, and 1 animal in experiment 3. Reactivation index was calculated as overlap divided by %eYFP+ and chance overlap was calculated as %eYFP*%cFos. Overlaps were normalized to chance by dividing the proportion of overlaps (i.e., overlaps/DAPI*100) by this chance calculation.

## 3. RESULTS

### 3.1 Reactivation of dDG engrams tagged during the first cocaine exposure in CPP does not induce locomotion

Male and female c57BL/6 mice received bilateral injections of either ChR2 or eYFP targeting the dDG to genetically label hippocampal ensembles active during a first cocaine exposure **(Figure 1A).** The number of DAPI **(Figure 1B, left)** and eYFP (**Figure 1B, right)** positive (^+^) labeled neurons were consistent across groups tagging approximately 7% of dDG neurons (*ns*, 3-way ANOVA). We have previously demonstrated effective DOX-dependent regulation of neuronal tagging using this specific TetTag viral prep (Kim et al., 2026). To further confirm DOX-dependent suppression of transgene expression, we additionally examined dDG tissue from mice maintained on DOX (n=5). Immunohistochemical analysis revealed c-Fos+ neurons with minimal detectable eYFP expression **(Figure 1C),** consistent with effective suppression of activity-dependent neuronal tagging under ON-DOX conditions.

**Figure 1.**
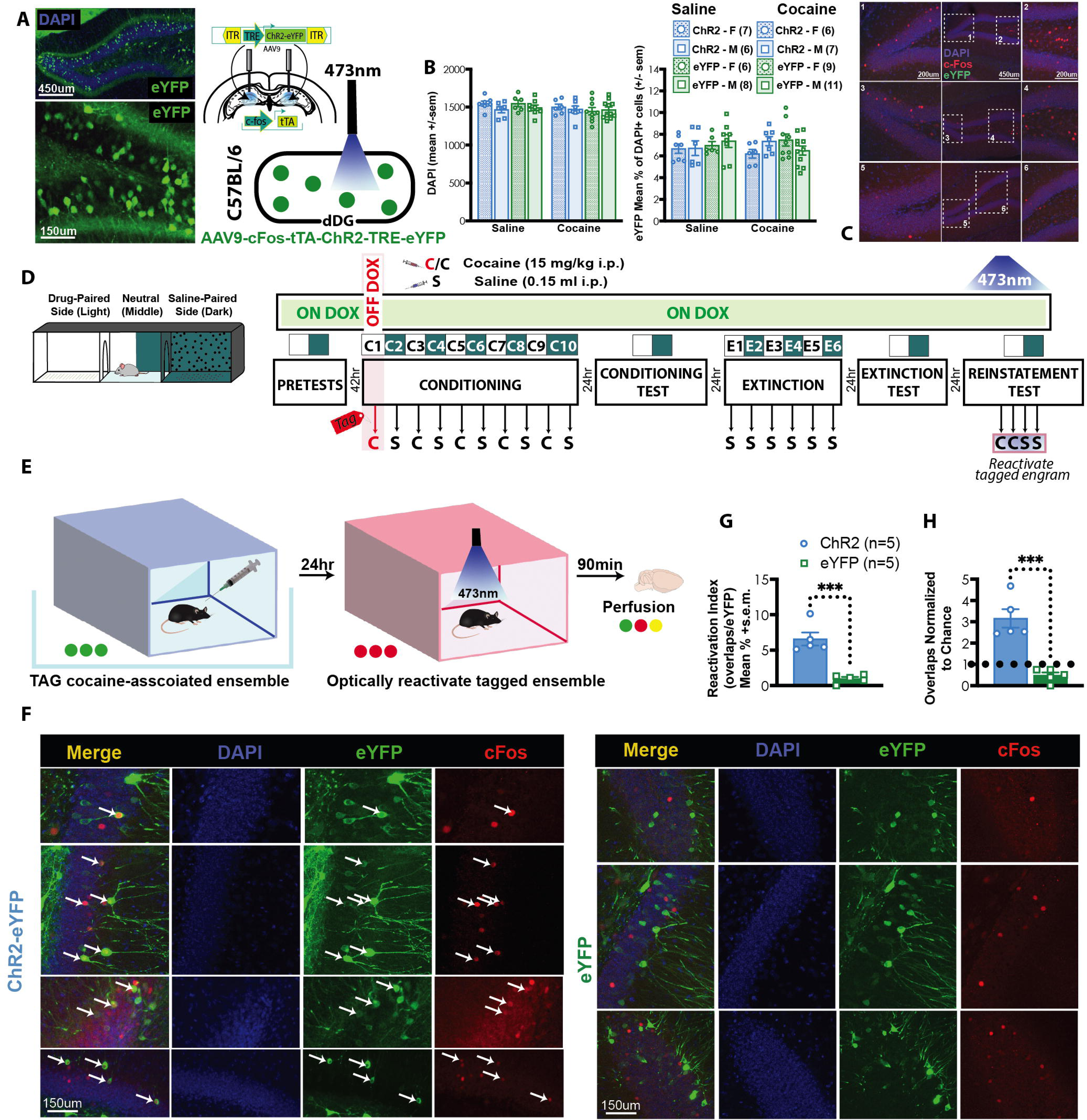
Experimental strategy and validation of activity-dependent neuronal tagging and optogenetic reactivation in the dorsal dentate gyrus. (A) Viral strategy and representative image for tagging hippocampal ensembles active during a first cocaine exposure in the dorsal dentate gyrus (dDG) using a TetTag strategy. (B) Quantification of DAPI+ and eYFP+ cells in the dDG. Approximately 7% of dDG neurons were tagged across groups. (C) Representative confocal images of the dorsal dentate gyrus from mice maintained on DOX (n=5). Sections were labeled for DAPI (blue), c-Fos (red), and eYFP (green). Enlarged regions illustrate the presence of c-Fos+ cells with minimal detectable eYFP expression under ON-DOX conditions. Scale bars: 450 μm (overview) and 150 μm (enlargements). (D) Apparatus and experimental timeline for cocaine conditioned place preference (CPP), extinction, and reinstatement procedure. (E) Experimental design and (F) representative fluorescence images of the dDG from ChR2-eYFP and eYFP-only control mice following optical stimulation. Neuronal ensembles tagged during cocaine exposure. Mice were subsequently placed in a novel context and received identical optical stimulation, followed by perfusion 90 min later for c-Fos immunohistochemistry. Images show eYFP-labeled neurons (green), c-Fos immunoreactivity (red), and merged channels (yellow). Overlapping cells are denoted with white arrows (G) Quantification of the percentage of eYFP+ neurons co-expressing c-Fos in ChR2-eYFP and eYFP-only mice following optical stimulation using reactivation index. Each point represents an individual animal. Data are presented as mean ± s.e.m. *p < 0.05, **p < 0.01, ***p < 0.001, ****p < 0.0001.

Following tagging, mice underwent cocaine CPP, extinction, and reinstatement procedures **(Figure 1D).** To determine whether reactivation of the tagged ensemble could influence reinstatement, all mice received optical stimulation during the reinstatement test, either alone or in combination with a cocaine priming injection. To verify that optical stimulation could effectively reactivate tagged neurons via ChR2, we conducted an additional experiment in which mice expressing either ChR2-eYFP or eYFP alone underwent tagging during a cocaine experience. The following day, they received optical stimulation and timed perfusions (90 min later) to assess neuronal activation. Stimulation was delivered in a distinct environment to distinguish context-dependent neuronal recruitment from optogenetically induced reactivation **(Figure 1E).** We found significantly greater c-Fos co-expression in eYFP-positive neurons in ChR2-eYFP mice compared with eYFP-only controls **(Figure 1F).** We demonstrated this using an ensemble-specific reactivation index (t_8_ = 5.851, p=0.0004; unpaired t-test) **(Figure 1F).** We also demonstrated this using a measure of overlaps normalized to chance [ME: GROUP; F_2,12_ = 21.29, p<0.0001; 1-way ANOVA) (ChR2 *vs.* chance: p=0.0002; eYFP *vs.* chance: p=3073) **(Figure 1G).**

#### 3.1.1. Conditioning

During conditioning, cocaine produced robust locomotor activation and sensitization developed across repeated drug exposures. Specifically, mice traveled more distance **(Figure 2A, left)** (Main Effect [ME] of TIME; F_4,280_ = 11.34, p<0.0001; 3-way Repeated Measures [RM] ANOVA), at higher speeds **(Figure 2B, left)** (ME:TIME; F_4,280_ = 3.6, p=0.0069; 3-way RM ANOVA), across days when cocaine was administered. These locomotor effects were not observed on saline days **(Figures 2A,B, right)** (*ns*, 3-way RM ANOVA). This was also evident when we collapsed data across days **(Figures 2C,D) (**Distance: SEX x DRUG INTERACTION; F_1,112_ = 4.883, p=0.0292; Speed: ME:DRUG; F_1,112_ = 248.7, p<0.0001; 3-way RM ANOVA), and when data were collapsed across sex **(Figures 2E-H)** (Figure 2E left: ME:TIME; F_4,232_ = 21.47, p<0.0001; Figure 2F left: ME:TIME; F_4,232_ = 7.004, p<0.0001; Figure 2G: DRUG x VIRUS INTERACTION; F_1,58_ = 5.477, p<0.0227; Figure 2H: ME:DRUG; F_1,58_ = 247, p<0.0001; 2-way ANOVA).

**Figure 2.**
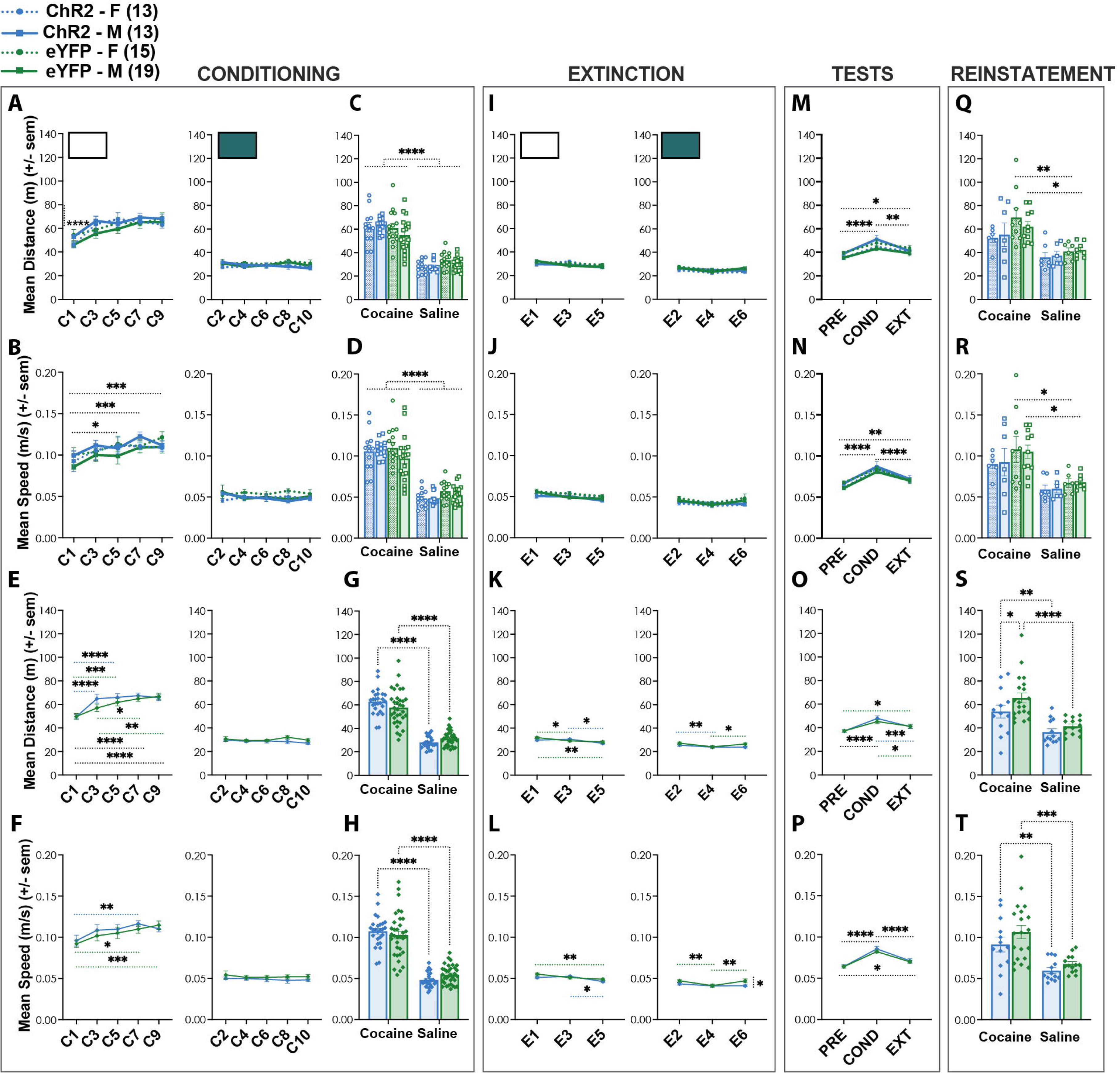
Reactivation of dDG engrams tagged during a first cocaine exposure does not induce locomotion during reinstatement. (A-H) Cocaine increased locomotion during conditioning, reflected by greater distance traveled and speed across cocaine sessions relative to saline sessions, demonstrating locomotor sensitization. (I-L) During extinction, locomotor activity progressively decreased across sessions. (M-P) Mice exhibited greater locomotion during the conditioning test despite testing occurring in a drug-free state. (Q-T) During reinstatement, optical reactivation of first-exposure cocaine-associated ensembles did not independently induce locomotion nor disrupt cocaine-induced locomotor activation. Cocaine increased both distance traveled and speed, with females showing greater sensitivity to locomotor-inducing effects. Data are presented as mean ± s.e.m. *p < 0.05, **p < 0.01, ***p < 0.001, ****p < 0.0001.

#### 3.1.2. Extinction

During extinction, we saw no group differences. Distance traveled **(Figure 2I)** (*ns*, 3-way RM ANOVA) and speed **(Figure 2J)** (*ns*, 3-way RM ANOVA) were similar to saline conditioning sessions. When collapsed across sex, we saw that mice traveled less distance **(Figure 2K)** (Figure 2K left: ME:TIME; F_2,116_ = 8.292, p=0.0004; Figure 2K right: ME:TIME; F_2,116_ = 5.855, p=0.0038; 2-way ANOVA) and moved slower **(Figure 2L)** (Figure 2L left: ME:TIME; F_2,116_ = 7.854, p=0.0006; Figure 2L right: ME:TIME; F_2,116_ = 5.095, p=0.0076; 2-way ANOVA) across sessions, except the last session (E6) where mice in the eYFP group traversed the apparatus at a higher speed than ChR2 mice (p=0.0185).

#### 3.1.3. Tests

Mice underwent three tests prior to reinstatement: a pre-test (PRE) to assess initial side preference, a post-conditioning test (COND) to evaluate the strength of CPP, and an extinction test (EXT) to determine whether preference for the drug-paired side was eliminated following extinction training. Although all tests were conducted in a drug-free state, mice exhibited greater locomotor activity, reflected by increased distance traveled **(Figure 2M) (**ME:TIME; F_2,168_ = 18.60, p<0.0001; 3-way RM ANOVA) and speed **(Figure 2N) (**ME:TIME; F_2,168_ = 43.85, p<0.0001; 3-way RM ANOVA), during the conditioning test. These effects were even more robust when we collapsed across sex **(Figures 2O, P)** (Distance: ME:TIME; F_2,116_ = 29.94, p<0.0001; Speed: ME:TIME; F_2,116_ = 63.79, p<0.0001; 2-way RM ANOVA).

#### 3.1.4. Reinstatement

During reinstatement, all mice received optical stimulation. Artificial reactivation of dDG engrams tagged during the first cocaine exposure did not independently induce locomotion nor did it disrupt cocaine-induced locomotor activation. As expected, mice administered cocaine exhibited greater distance traveled **(Figure 2Q)** (ME:DRUG; F_1,52_ = 25.14, p<0.0001; ME:SEX, F_1,52_ = 4.173, p=0.0462, 3-way ANOVA) and higher locomotor speeds **(Figure 2R)** (ME:DRUG; F_1,52_ = 22.50, p<0.0001; 3-way ANOVA) compared to saline treated mice. Interestingly the magnitude of this cocaine-induced increase was greater in eYFP controls than in ChR2 mice and females were more sensitive (p=0.0039). This effect became more apparent when data were collapsed across sex, revealing a significant difference between ChR2 and eYFP mice (p=0.0397) in the cocaine condition for both distance traveled **(Figure 2S)** (ME:DRUG; F_1,56_ = 26.24, p<0.0001; ME:VIRUS, F_1,52_ = 4.254, p=0.0438, 3-way ANOVA) and speed **(Figure 2T)** (ME:DRUG; F_1,56_ = 24.38, p<0.0001; 3-way ANOVA).

### 3.2 Reactivation of dDG engrams tagged during the first cocaine exposure attenuates cocaine-primed reinstatement

We measured preference in three ways. First, we calculated a *preference ratio* showing a comparison of time spent on each side of the apparatus (excluding the middle). All mice initially preferred the dark side **(Figure 3A)** (ME:SIDE, F_1,112_ = 423.3, p<0.0001, 3-way RM ANOVA), therefore cocaine was paired with the light side. Following conditioning, mice preferred this side **(Figure 3B)** (ME:SIDE, F_1,112_ = 276.9, p<0.0001, 3-way RM ANOVA). This preference was then extinguished until mice again preferred the dark side by replacing cocaine with saline **(Figure 3C)** (ME:SIDE, F_1,112_ = 298.3, p<0.0001, 3-way RM ANOVA). These effects were also evident when collapsed across sex **(Figures 3D-F)** (Figure 3D: ME:SIDE, F_1,58_ = 216, p<0.0001; Figure 3E: ME:SIDE, F_1,56_ = 145, p<0.0001; Figure 3F: ME:SIDE, F_1,56_ = 151.4, p<0.0001; 2-way RM ANOVA).

**Figure 3.**
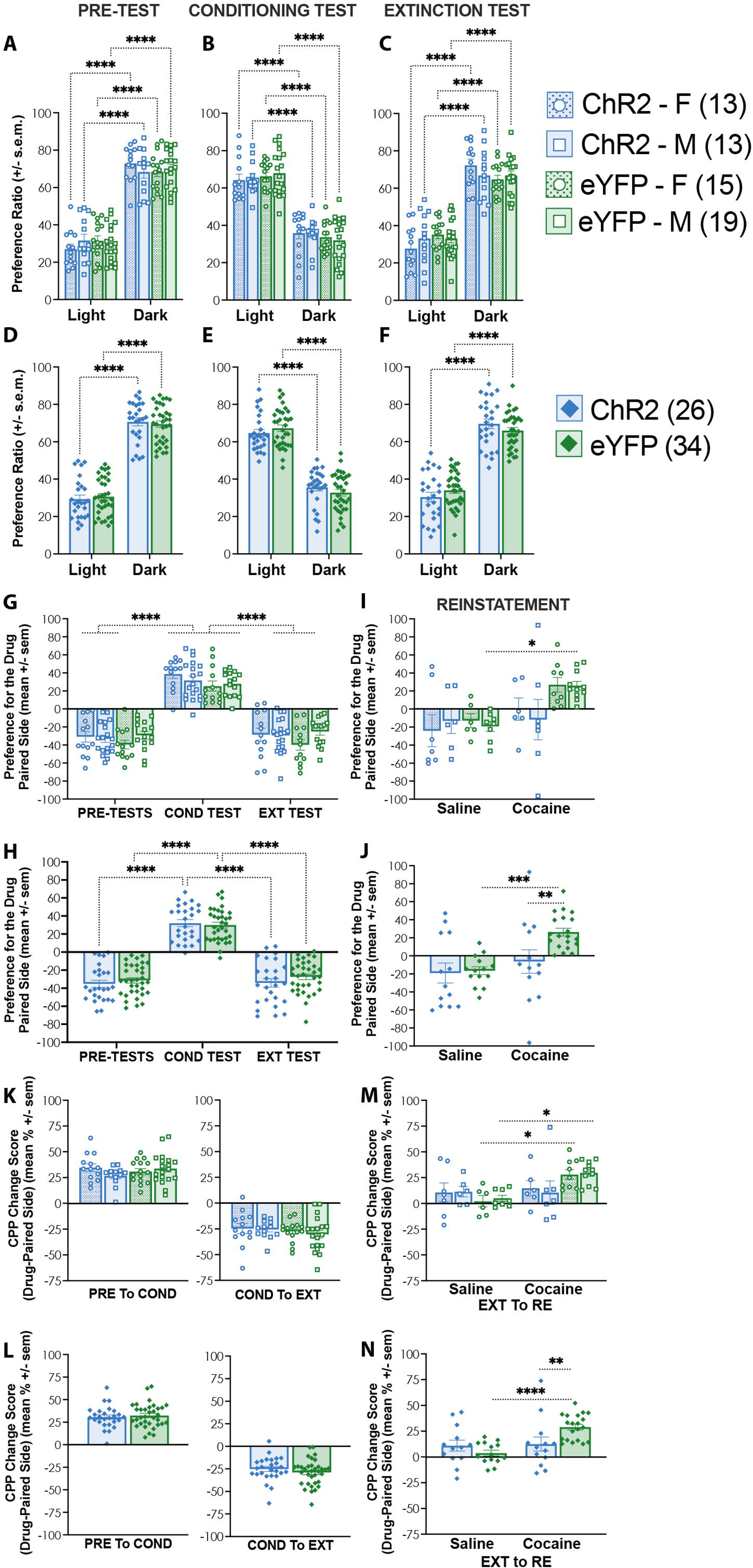
Reactivation of dDG engrams tagged during a first cocaine exposure attenuates cocaine-primed reinstatement. (A–F) Preference ratio across pre-test, conditioning, and extinction sessions. Mice initially preferred the dark side, shifted preference to the cocaine-paired light side following conditioning, and returned to preferring the dark side after extinction. (G,H) Preference scores across tests demonstrating acquisition and extinction of CPP. (I,J) During reinstatement, cocaine priming reinstated CPP in eYFP mice. Optical reactivation of first-exposure cocaine-associated ensembles did not independently reinstate CPP but attenuated cocaine-induced reinstatement relative to cocaine-primed eYFP controls. (K–N) CPP change scores across testing phases. No group differences emerged from pre-test to conditioning or conditioning to extinction. During reinstatement, cocaine increased CPP in eYFP mice, whereas ChR2 mice showed reduced cocaine-induced reinstatement. Data are presented as mean ± s.e.m. *p < 0.05, **p < 0.01, ***p < 0.001, ****p < 0.0001.

We then calculated a *preference score,* specifically for the drug paired side, to show that preference changed across pre-tests to conditioning to extinction **(Figure 3G)** (ME:TEST, F_2,112_ = 234.6, p<0.0001, 3-way RM ANOVA) and this was also seen collapsed across sex **(Figure 3H)** (ME:TEST, F_2,116_ = 243.5, p<0.0001; 2-way RM ANOVA). During reinstatement, all mice were given optical stimulation and parsed into groups that either received an injection of cocaine to reinstate drug-seeking or a saline control **(Figure 1D).** In eYFP groups, a priming injection of cocaine successfully reinstated drug-seeking **(Figure 3I)** (ME:DRUG, F_1,31_ = 10.04, p=0.0034, 3-way ANOVA). When collapsed across sex (ME:VIRUS, F_1,56_ = 4.503, p=0.0383; ME:DRUG, F_1,56_ = 11.22, p=0.0015; 2-way ANOVA), we saw a significant difference between saline and cocaine treatment in the eYFP group (p=0.0003) **(Figure 3J**). Importantly, reactivation of hippocampal ensembles encoding a first cocaine exposure altered cocaine-seeking behavior during the reinstatement test **(Figures 3I,J).** Optical stimulation of dDG ensembles tagged during the initial cocaine experience did not independently reinstate CPP (*ns*, 3-way ANOVA), indicating that artificial reactivation of the cocaine-related memory was insufficient to substitute for cocaine priming **(Figure 3I,J)**. However, when optical reactivation was paired with a cocaine priming injection, mice showed reduced reinstatement relative to cocaine-primed controls (p=0.0053) **(Figure 3J**), suggesting that reactivation of the early cocaine memory interfered with cocaine-induced drug-seeking.

Finally, we calculated a *CPP change score,* showing the change in preference for the drug-paired side across tests **(Figures 3K-N).** There were no group differences in this score from pre-test to the conditioning test **(Figures 3K,L** **left)** (*ns*, 3-way ANOVA, 2-way ANOVA) or from the conditioning test to the extinction test **(Figures 3K,L** **right)** (*ns*, 3-way ANOVA, 2-way ANOVA). At reinstatement, group differences emerged between saline and cocaine conditions driven primarily by eYFP mice **(Figures 3M,N)** (DRUG x VIRUS INTERACTION, F_1,52_ = 6.641, p=0.0128; 3-way ANOVA) showing significant differences between saline- and cocaine-treated groups in both females (p=0.0262) and males (p=0.0142) whereas ChR2 mice did not show this effect **(Figure 3M).** When sex was collapsed **(Figure 3N),** the same pattern remained evident (DRUG x VIRUS INTERACTION, F_1,56_ = 7.187, p=0.0096; 2-way ANOVA). Post hoc analyses revealed a significant difference between saline- and cocaine-treated eYFP mice (p<0.0001) and within the cocaine condition, ChR2 mice differed significantly from eYFP controls (p=0.0081).

### 3.3 Reactivation of dDG engrams tagged during the first and fourth cocaine exposure in the open field induces context-dependent locomotion in females

To determine whether these effects generalized beyond CPP, we examined locomotor activity in an open field assay. Because locomotor sensitization is highly context-dependent (Mattson et al., 2008), we tested whether reactivation of dDG cocaine-associated ensembles could induce locomotor responses when reactivation occurred in the original encoding environment. Ensembles active during either the first or fourth cocaine exposure were reactivated during a drug-free test session (only NO-VIRUS mice received cocaine as a positive control for locomotor sensitization), and we examined whether locomotor responses differed as a function of exposure history and sex **(Figure 4A).**

**Figure 4.**
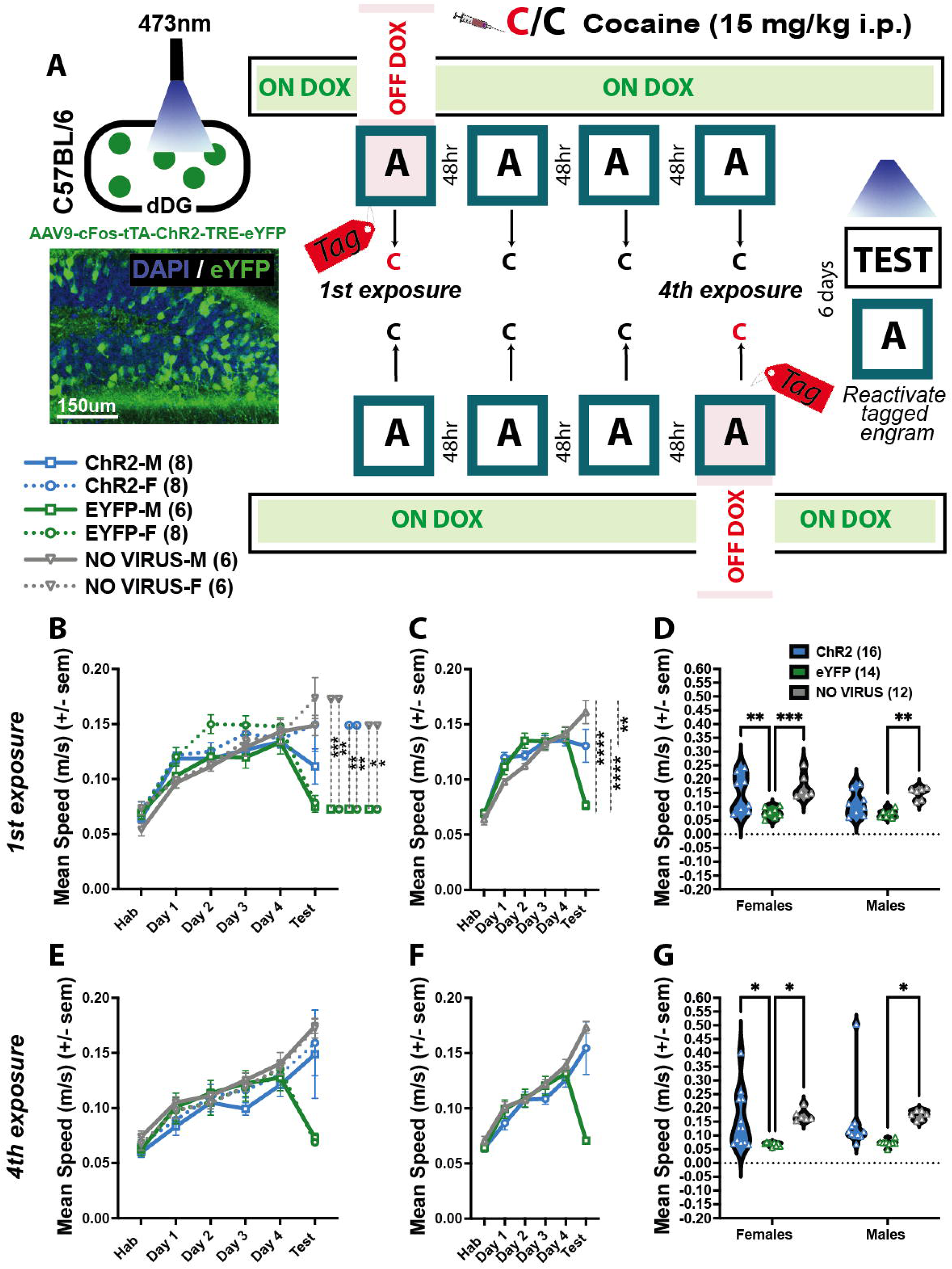
Reactivation of dDG cocaine-associated ensembles in the open field induces context-dependent locomotor effects in females. (A) Experimental design for open field-testing following tagging of either first- or fourth-exposure cocaine ensembles. Mice received repeated cocaine exposures in the open field and underwent drug-free optical reactivation testing. (B–D) First cocaine exposure tagged. Cocaine increased locomotion across conditioning sessions and sensitization developed over time. Reactivation of first-exposure cocaine-associated ensembles induced locomotor effects selectively in females. (E–G) Fourth cocaine exposure tagged. Cocaine again increased locomotion across conditioning sessions. Reactivation of fourth-exposure cocaine-associated ensembles induced locomotor activity that more closely resembled cocaine-induced sensitization, particularly in females. Locomotor effects were dissociated from CPP behavior. Data are presented as mean ± s.e.m. *p < 0.05, **p < 0.01, ***p < 0.001, ****p < 0.0001.

#### 3.3.1 First exposure

Cocaine increased locomotion across conditioning sessions and sensitization developed from the first to the final cocaine exposure **(Figures 4B)** (DAY x VIRUS INTERACTION, F_10,190_ = 8.132, p<0.0001; 3-way RM ANOVA), an effect that remained when data were collapsed across sex **(Figure 4C)** (DAY x VIRUS INTERACTION, F_10,246_ = 8.522, p<0.0001; 2-way RM ANOVA). Reactivation of first-exposure cocaine ensembles did not induce locomotor activation in males, however, females showed greater sensitivity to the locomotor-inducing effects of cocaine memory reactivation. On test day, significant differences were observed between female eYFP and ChR2 groups (p=0.008) and between female eYFP and No-Virus groups (p=0.0012) **(Figure 4B)**. When collapsed across sex, the interaction was driven by test day differences between eYFP and No Virus groups (p<0.0001), eYFP and ChR2 (p<0.0001) and ChR2 and No-Virus (p=0.006) suggesting that reactivation of first exposure cocaine ensembles induced locomotor activity, although not to the extent produced by cocaine itself. Looking specifically at the test session **(Figure 4D),** we saw a significant main effect of virus (F_1,38_ = 3.146, p<0.0001; 2-way ANOVA). Post hoc analyses revealed significant difference between No-Virus and eYFP groups in both males (p=0.0053) and females (p=0.0003), as expected. Importantly, females also showed a significant difference between ChR2 and eYFP groups (p=0.0037).

#### 3.3.2 Fourth exposure

Similarly, cocaine increased locomotion across conditioning sessions and sensitization developed from the first to the final cocaine exposure **(Figures 4E)** (DAY x VIRUS INTERACTION, F_10,225_ = 3.423, p=0.00034; 3-way RM ANOVA), an effect that remained when data were collapsed across sex **(Figure 4F)** (DAY x VIRUS INTERACTION, F_10,240_ = 6.557, p<0.0001; 2-way RM ANOVA). Here, reactivation of fourth-exposure cocaine ensembles produced more comparable locomotor effects across males and females, although females still appeared more sensitive to the effects of memory reactivation. When collapsed across sex, the interaction was driven by test day differences between eYFP and No Virus groups (p<0.0001), eYFP and ChR2 (p<0.0001). Unlike the first-exposure experiment, there was no significant difference between ChR2 and No-Virus groups, suggesting that reactivation of fourth-exposure cocaine ensembles induced locomotor activity to a level comparable to cocaine. Focusing specifically on locomotor behavior during the test session **(Figure 4G),** we saw a significant main effect of virus (F_2,45_ = 7.965, p=0.0011; 2-way ANOVA). As expected, eYFP mice differed significantly from No-Virus controls in both males (p = 0.0480) and females (p = 0.0368). Notably, females also showed a significant difference between ChR2 and eYFP groups (p = 0.0368), further supporting greater sensitivity to cocaine memory reactivation in females. Importantly, these locomotor effects were dissociated from place preference, indicating that the attenuation of reinstatement was not driven by nonspecific changes in activity.

### 3.4 Reactivation of dDG engrams tagged during the fourth cocaine exposure in CPP induce locomotion in females

We next asked whether a similar attenuation of cocaine-primed reinstatement would be observed when the reactivated dDG ensemble was tagged during the fourth, rather than the first, cocaine exposure. Here, the experimental design was the same as experiment 1, except that we labeled hippocampal ensembles associated with the fourth cocaine exposure **(Figure 5A).** The number of DAPI^+^ **(Figure 5B, left)** and eYFP^+^ (**Figure 5B, right)** cells were consistent across groups, once again tagging approximately 7% of dDG neurons (*ns*, 3-way ANOVA). Following conditioning day C6, mice were taken off-DOX and conditioning day C7 was tagged 42 hours later **(Figure 5C).** All mice received optical stimulation during the reinstatement test: either alone or in combination with a cocaine priming injection.

**Figure 5.**
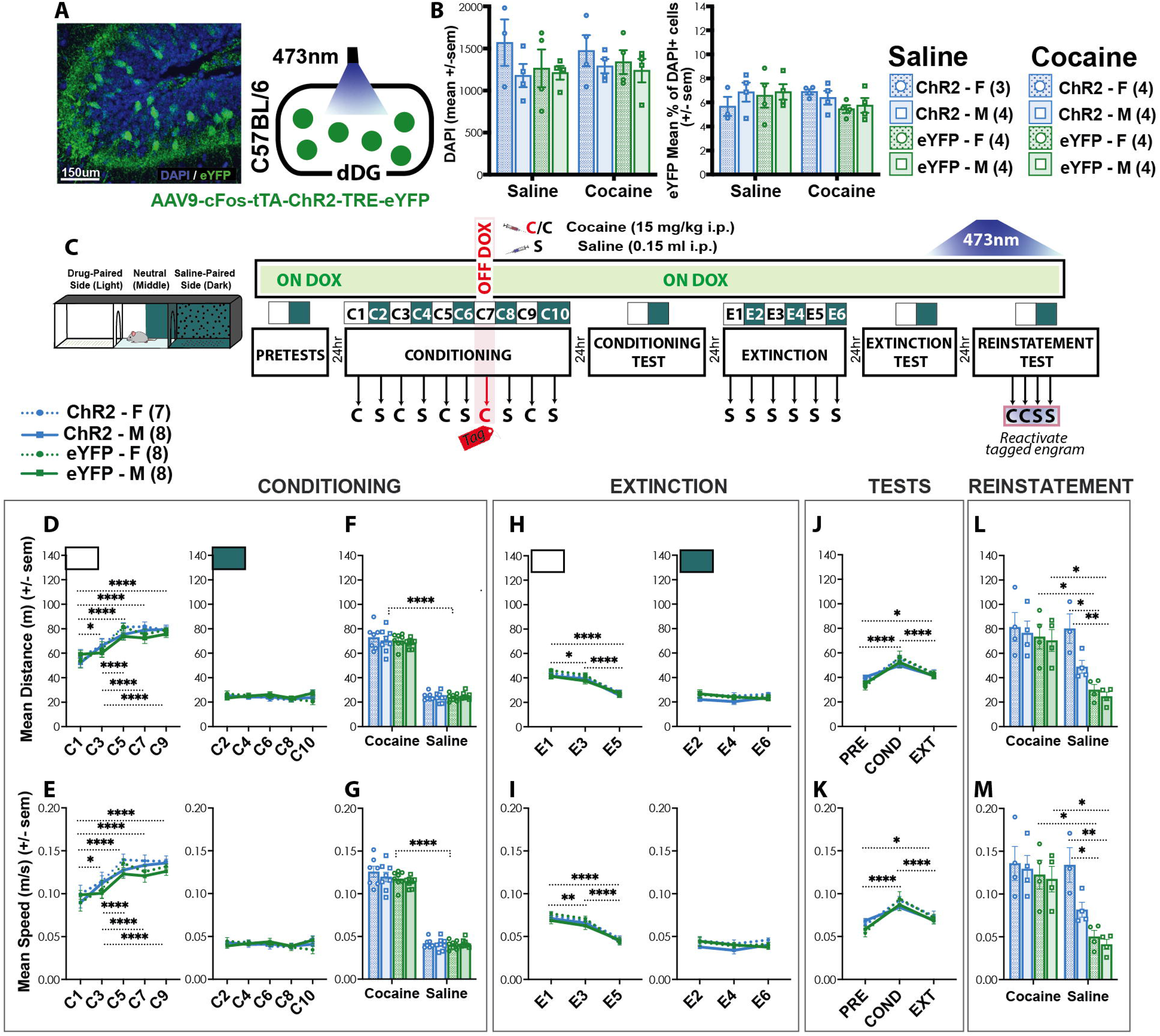
Reactivation of dDG engrams tagged during a fourth cocaine exposure induces locomotion in females during reinstatement. (A) Viral strategy and representative image for tagging hippocampal ensembles active during a fourth cocaine exposure in the dorsal dentate gyrus (dDG) using a TetTag strategy. (B) Quantification of DAPI+ and eYFP+ cells in the dDG. Approximately 7% of dDG neurons were tagged across groups. (C) Apparatus and experimental timeline for cocaine conditioned place preference (CPP), extinction, and reinstatement procedure. (D–G) Cocaine increased locomotion during conditioning and sensitization developed across repeated exposures. (H,I) Locomotor activity decreased during extinction. (J,K) Increased locomotion was observed during the conditioning test despite the absence of cocaine. (L,M) During reinstatement, optical reactivation of fourth-exposure cocaine-associated ensembles induced locomotor activity in females but did not disrupt cocaine-induced locomotion. Saline-treated female ChR2 mice displayed locomotor activity comparable to cocaine-treated female ChR2 mice, indicating enhanced sensitivity to cocaine memory reactivation in females. Data are presented as mean ± s.e.m. *p < 0.05, **p < 0.01, ***p < 0.001, ****p < 0.0001.

#### 3.4.1 Conditioning

As expected, cocaine produced locomotor activation and sensitization across drug sessions with mice traveling more distance **(Figure 5D, left)** (ME: TIME; F_4,108_ = 32.08, p<0.0001; 3-way RM ANOVA), and exhibiting higher speeds **(Figure 5E, left)** (ME:TIME; F_4,108_ = 31.22, p<0.0001; 3-way RM ANOVA). These locomotor effects were absent across saline days **(Figures 5D,E, right)** (*ns*, 3-way RM ANOVA). We saw this when we collapsed the data across days as well **(Figures 5F,G) (**Distance: ME: DRUG; F_1,27_ = 1168, p<0.0001; Speed: ME:DRUG; F_1,27_ = 1187, p<0.0001; 3-way RM ANOVA).

#### 3.4.2 Extinction

No group differences were observed during extinction. Both distance **(Figure 5H, left)** (ME:TIME; F_2,54_ = 124.1, p<0.0001; 3-way RM ANOVA) and speed **(Figure 5I, left)** ME:TIME; F_2,54_ = 126.7, p<0.0001; 3-way RM ANOVA) decreased across sessions in the formerly drug-paired context. During the sessions where saline was also previously administered there were no significant differences in distance traveled **(Figure 5H, right)** or speed **(Figure 5I, right)** (ns, 3-way RM ANOVA).

#### 3.4.3 Tests

As before, even though tests were conducted in a drug-free state, mice exhibited greater locomotor activity, reflected by increased distance traveled **(Figure 5J) (**ME:TIME; F_2,54_ = 36.50, p<0.0001; 3-way RM ANOVA) and speed **(Figure 5K) (**ME:TIME; F_2,54_ = 34.94, p<0.0001; 3-way RM ANOVA), during the conditioning test.

#### 3.4.4. Reinstatement

All mice received optical stimulation during the reinstatement test. Here, artificial reactivation of dDG engrams tagged during the fourth cocaine exposure did induce locomotor activity in females. However, it did not disrupt cocaine-induced locomotor activation **(Figures 5L,M)** (Distance: DRUG x VIRUS INTERACTION; F_1,23_ = 6.047, p=0.0219; Speed: DRUG x VIRUS INTERACTION; F_1,23_ = 5.933, p=0.023; 3-way ANOVA). For both distance traveled and speed, cocaine increased locomotion as expected. However, among saline-treated mice, ChR2 mice also showed elevated locomotor activity following memory reactivation. This increase was modest and not significant in males but was robust in females. In fact, saline-treated female ChR2 mice were indistinguishable from cocaine-treated female ChR2 mice for both distance traveled and speed. Saline-treated female ChR2 mice also differed significantly from saline-treated eYFP females (distance: p = 0.0152; speed: p = 0.0141) and saline-treated eYFP males (distance: p = 0.0057; speed: p = 0.0053).

### 3.5 Reactivation of dDG engrams tagged during the fourth cocaine exposure does not attenuate cocaine-primed reinstatement

As before, all mice initially preferred the dark side **(Figure 6A)** (ME:SIDE, F_1,27_ = 80.9, p<0.0001, 3-way RM ANOVA) thus, cocaine was paired with the light side. Following conditioning, preference switched to the light side **(Figure 6B)** (ME:SIDE, F_1,27_ = 75, p<0.0001, 3-way RM ANOVA). During extinction, cocaine was replaced with saline causing extinction of this preference and mice once again preferred the dark side **(Figure 6C)** (ME:SIDE, F_1,27_ = 55.6, p<0.0001, 3-way RM ANOVA). These effects were also evident when collapsed across sex **(Figures 6D-F)** (Figure 6D: ME:SIDE, F_1,29_ = 85.18, p<0.0001; Figure 6E: ME:SIDE, F_1,29_ = 79.84, p<0.0001; Figure 6F: ME:SIDE, F_1,29_ = 58.53, p<0.0001; 2-way RM ANOVA).

**Figure 6.**
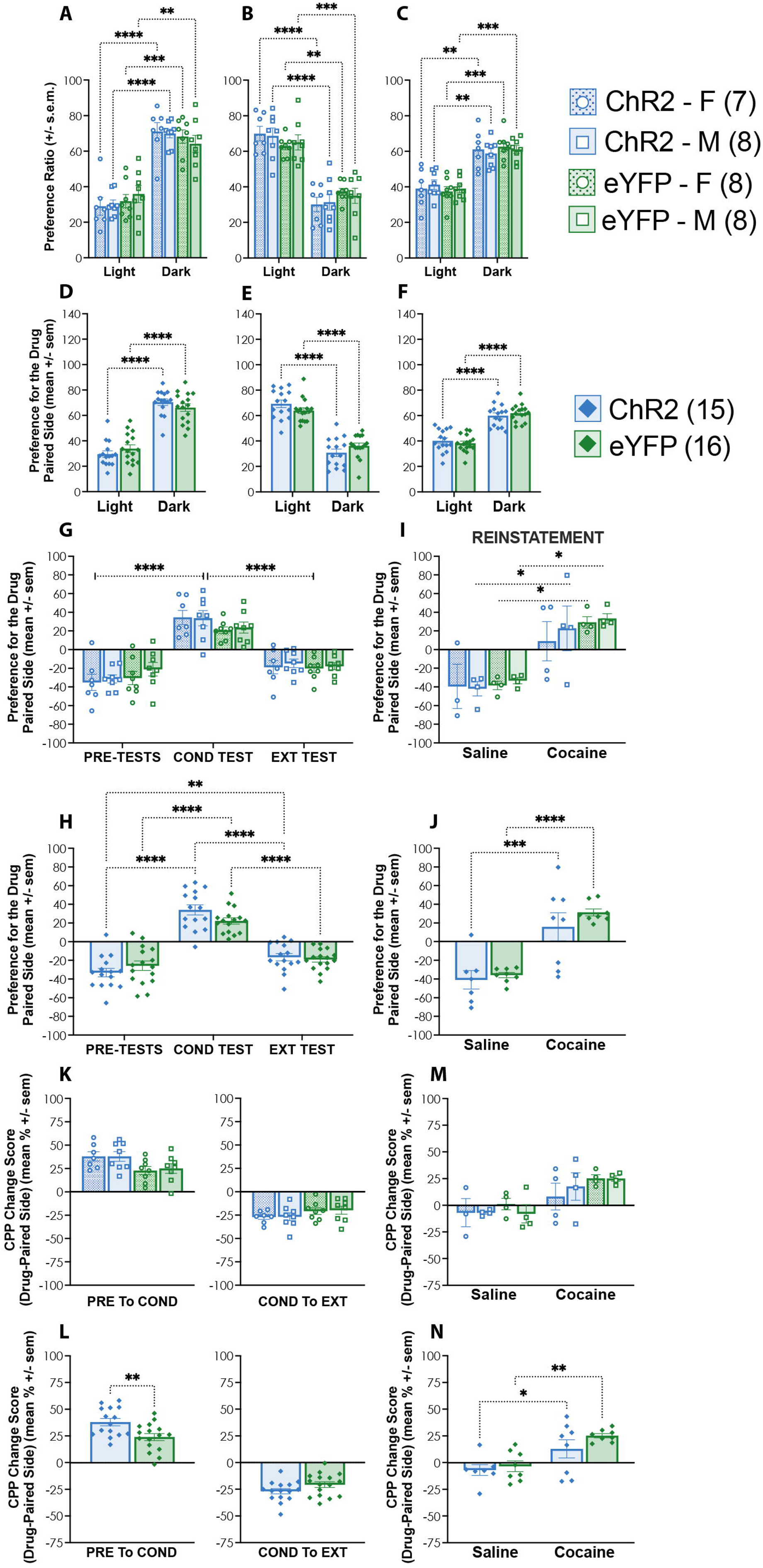
Reactivation of dDG engrams tagged during a fourth cocaine exposure does not attenuate cocaine-primed reinstatement. (A–F) Preference ratio across pre-test, conditioning, and extinction sessions. Mice initially preferred the dark side, shifted preference to the cocaine-paired light side following conditioning, and returned to preferring the dark side after extinction. (G,H) Preference scores across testing phases demonstrating acquisition and extinction of CPP. ChR2 mice showed less recovery of their original dark-side preference during extinction compared to eYFP controls. (I,J) Cocaine priming reinstated CPP in both eYFP and ChR2 groups. In contrast to first-exposure ensemble reactivation, reactivation of fourth-exposure cocaine-associated ensembles did not attenuate cocaine-induced reinstatement. (K–N) CPP change scores across testing phases. ChR2 mice showed stronger initial conditioning responses, but no group differences emerged during extinction. During reinstatement, cocaine increased CPP in both eYFP and ChR2 mice regardless of virus condition. Data are presented as mean ± s.e.m. *p < 0.05, **p < 0.01, ***p < 0.001, ****p < 0.0001.

**Figure 6G** further demonstrates how preference changed across testing (ME:TEST, F_2,54_ = 122.5, p<0.0001, 3-way RM ANOVA), which was also observed with data collapsed across sex **(Figure 6H)** (TEST x VIRUS INTERACTION, F_2,58_ = 3.282, p=0.0446; 2-way RM ANOVA). Here, ChR2 mice showed less recovery of their original preference for the dark side during extinction compared to eYFP mice as there was a significant difference between pre-tests and extinction in this group driving the interaction (p=0.0097). During reinstatement, a cocaine prime reinstated drug-seeking behavior **(Figure 6I)** (ME:DRUG, F_1,23_ = 38.30, p<0.0001; 3-way ANOVA). This effect was observed in both eYFP males (p = 0.0401) and females (p = 0.0364), and remained significant when data were collapsed across sex (ME:DRUG, F₁,₂₇ = 44.72, p < 0.0001; 2-way ANOVA), where eYFP mice given cocaine showed greater preference for the drug-paired side than saline controls (p < 0.0001) **(Figure 6J).** In contrast to reactivation of ensembles encoding a first cocaine exposure, reactivation of hippocampal ensembles encoding a fourth cocaine exposure did not reduce cocaine-seeking during reinstatement **(Figures 6I,J).** ChR2 mice given cocaine still showed a significant preference for the drug-paired side compared to saline-treated ChR2 mice, indicating that memory reactivation alone was insufficient to induce reinstatement. Although the effect appeared slightly attenuated in females, there were no significant sex differences within the ChR2 cocaine group **(Figure 6I).** Consistent with this, when data were collapsed across sex, only a main effect of drug was observed, with no effect of virus (ME:DRUG, F₁,₂₇ = 44.72, p < 0.0001; 2-way ANOVA).

**Figures 6K and 6L** show changes in preference for the drug-paired side across testing phases. The increase in CPP from the pre-tests to the conditioning test was greater in ChR2 mice, although this effect only emerged when we collapsed the data across sex (Figure 6K left: ns, 3-way RM ANOVA; Figure 6L left: t_29_, = 2.95, p < 0.0062; unpaired t-test). The stronger initial conditioning observed in ChR2 mice may have contributed to the reduced efficacy of memory reactivation in attenuating subsequent drug-seeking. There were no group differences in CPP from the conditioning test to the extinction test **(Figures 6K,L** **right)** (*ns*, 3-way ANOVA, 2-way ANOVA). During reinstatement, there was a main effect of drug, (ME:DRUG, F_1,23_ = 16.17, p=0.0005; 3-way ANOVA) **(Figure 6M).** When data were collapsed across sex (ME:DRUG, F_1,27_ = 18.13, p=0.0002; 2-way ANOVA) **(Figure 6N)** a main effect of drug remained, with both eYFP (p=0.0012) and ChR2 (p=0.0221) mice showing greater CPP change scores following cocaine compared to saline treatment.

### 3.6 Reactivation of dDG engrams tagged during the first saline exposure in CPP does not induce locomotion

Male and female c57BL/6 mice received bilateral injections of ChR2 or eYFP in the dDG to label hippocampal ensembles active during a first saline exposure **(Figure 7A).** The number of DAPI^+^ **(Figure 7B, left)** and eYFP^+^ (**Figure 7B, right)** labeled neurons were consistent across groups tagging approximately 9% of dDG neurons (*ns*, 3-way ANOVA), which was slightly more cells than the cocaine-associated ensembles that were tagged in the previous experiments **(Figure 7C).** The experimental set up was identical to experiment 1, except that on the first tagged conditioning day, mice were placed in the dark side and given saline **(Figure 7D).** To determine whether reactivation of the tagged ensemble could influence reinstatement, all mice received optical stimulation during reinstatement, either alone or in combination with a cocaine prime.

**Figure 7.**
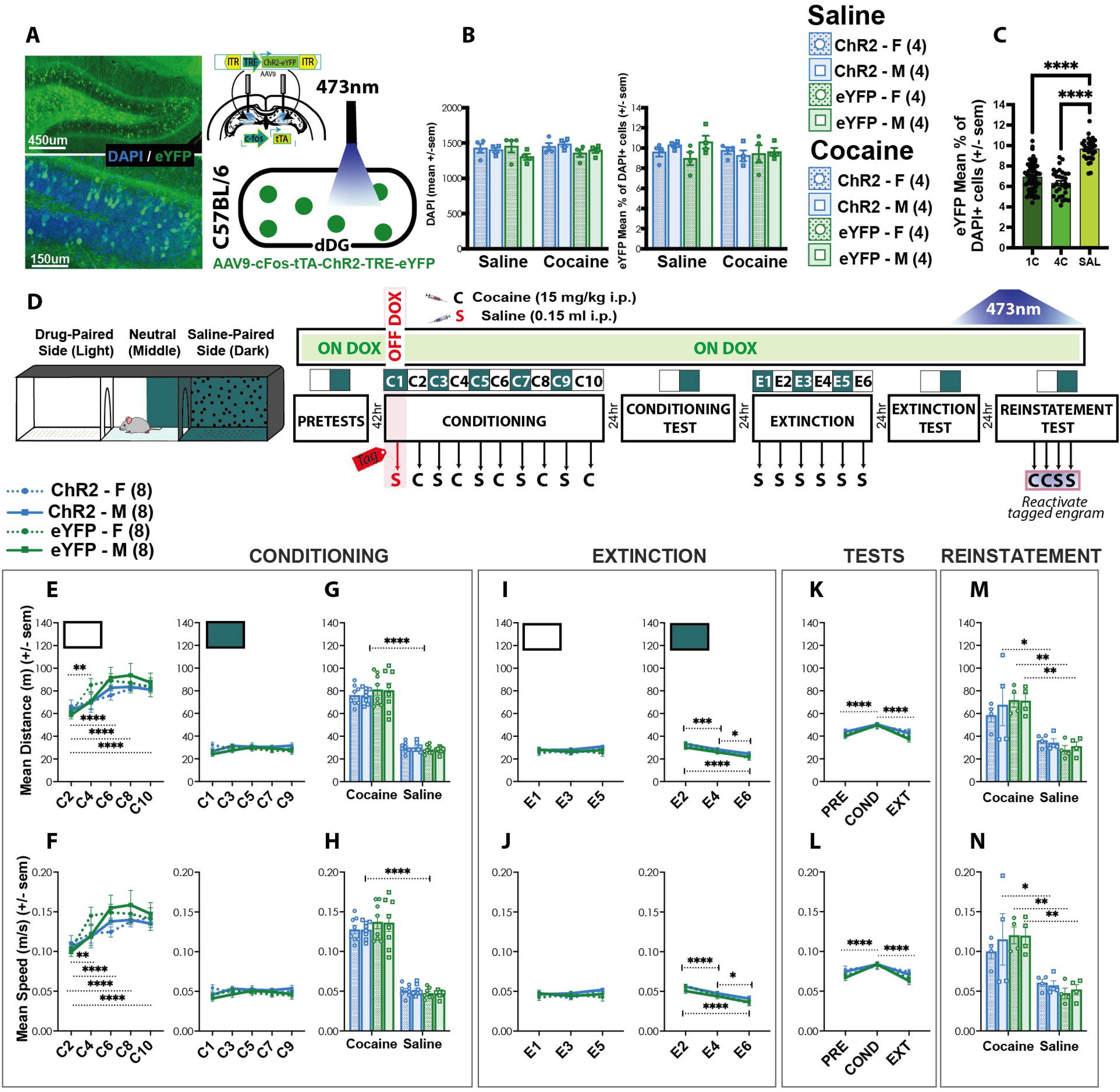
Reactivation of dDG engrams tagged during a first saline exposure does not induce locomotion during reinstatement. (A) Viral strategy and representative image for tagging hippocampal ensembles active during a fourth cocaine exposure in the dorsal dentate gyrus (dDG) using a TetTag strategy. (B) Quantification of DAPI+ and eYFP+ cells in the dDG. Approximately 9% of dDG neurons were tagged across groups. (C) Comparison of ensemble size across cocaine- and saline-tagged experiments. (D) Apparatus and experimental timeline for cocaine conditioned place preference (CPP), extinction, and reinstatement procedure. (E–H) Cocaine increased locomotion during conditioning and sensitization developed across repeated exposures. (I,J) Locomotor activity progressively decreased during extinction. (K,L) Greater locomotor activity was observed during the conditioning test relative to pre-test and extinction tests. (M,N) During reinstatement, optical reactivation of saline-tagged ensembles did not induce locomotion or disrupt cocaine-induced locomotor activation. Data are presented as mean ± s.e.m. *p < 0.05, **p < 0.01, ***p < 0.001, ****p < 0.0001.

#### 3.6.1 Conditioning

As before, cocaine produced locomotor activation and sensitization across drug sessions in terms of distance **(Figure 7E, left)** (ME: TIME; F_4,112_ = 17.18, p<0.0001; 3-way RM ANOVA), and speed **(Figure 7F, left)** (ME:TIME; F_4,112_ = 16.53, p<0.0001; 3-way RM ANOVA). These locomotor effects were not seen across saline days **(Figures 7E,F, right)** (*ns*, 3-way RM ANOVA). This was observed when we collapsed the data across days also **(Figures 7G,H) (**Distance: ME: DRUG; F_1,28_ = 627.2, p<0.0001; Speed: ME:DRUG; F_1,27_ = 586.5, p<0.0001; 3-way RM ANOVA).

#### 3.6.2 Extinction

No group differences were observed during extinction. During sessions where saline was previously administered there were no significant differences in distance traveled **(Figure 7I, left)** or speed **(Figure 7J, left)** (ns, 3-way RM ANOVA). Both distance **(Figure 7I, right)** (ME:TIME; F_2,56_ = 27.59, p<0.0001; 3-way RM ANOVA) and speed **(Figure 7J, right)** ME:TIME; F_2,56_ = 29.81, p<0.0001; 3-way RM ANOVA) decreased across sessions in the formerly drug-paired context.

#### 3.6.3 Tests

Mice exhibited greater locomotor activity, reflected by increased distance traveled **(Figure 7K) (**ME:TIME; F_2,56_ = 16.71, p<0.0001; 3-way RM ANOVA) and speed **(Figure 7L) (**ME:TIME; F_2,56_ = 15.33, p<0.0001; 3-way RM ANOVA), during the conditioning test.

#### 3.6.4 Reinstatement

Artificial reactivation of dDG engrams tagged during the first saline exposure did not induce locomotion nor did it disrupt cocaine-induced locomotor activation. As expected, mice administered cocaine exhibited greater distance traveled **(Figure 7M)** (ME:DRUG; F_1,24_ = 39.17, p<0.0001; 3-way ANOVA) and higher locomotor speeds **(Figure 7N)** (ME:DRUG; F_1,24_ = 37.61, p<0.0001; 3-way ANOVA) compared to saline treated mice. **Reactivation of dDG engrams tagged during the first saline exposure attenuates cocaine-primed reinstatement in females**

All mice initially preferred the dark side **(Figure 8A)** (ME:SIDE, F_1,28_ = 96.69, p<0.0001, 3-way RM ANOVA), so cocaine was again paired with the light side. Following conditioning, preference switched to the light side **(Figure 8B)** (ME:SIDE, F_1,28_ = 45.68, p<0.0001, 3-way RM ANOVA). Following extinction, mice again preferred the dark side **(Figure 8C)** (ME:SIDE, F_1,28_ = 90.23, p<0.0001, 3-way RM ANOVA). These effects were also evident when collapsed across sex **(Figures 8D-F)** (Figure 8D: ME:SIDE, F_1,30_ = 99.37, p<0.0001; Figure 8E: ME:SIDE, F_1,30_ = 48.26, p<0.0001; Figure 8F: ME:SIDE, F_1,29_ = 87.76, p<0.0001; 2-way RM ANOVA).

**Figure 8.**
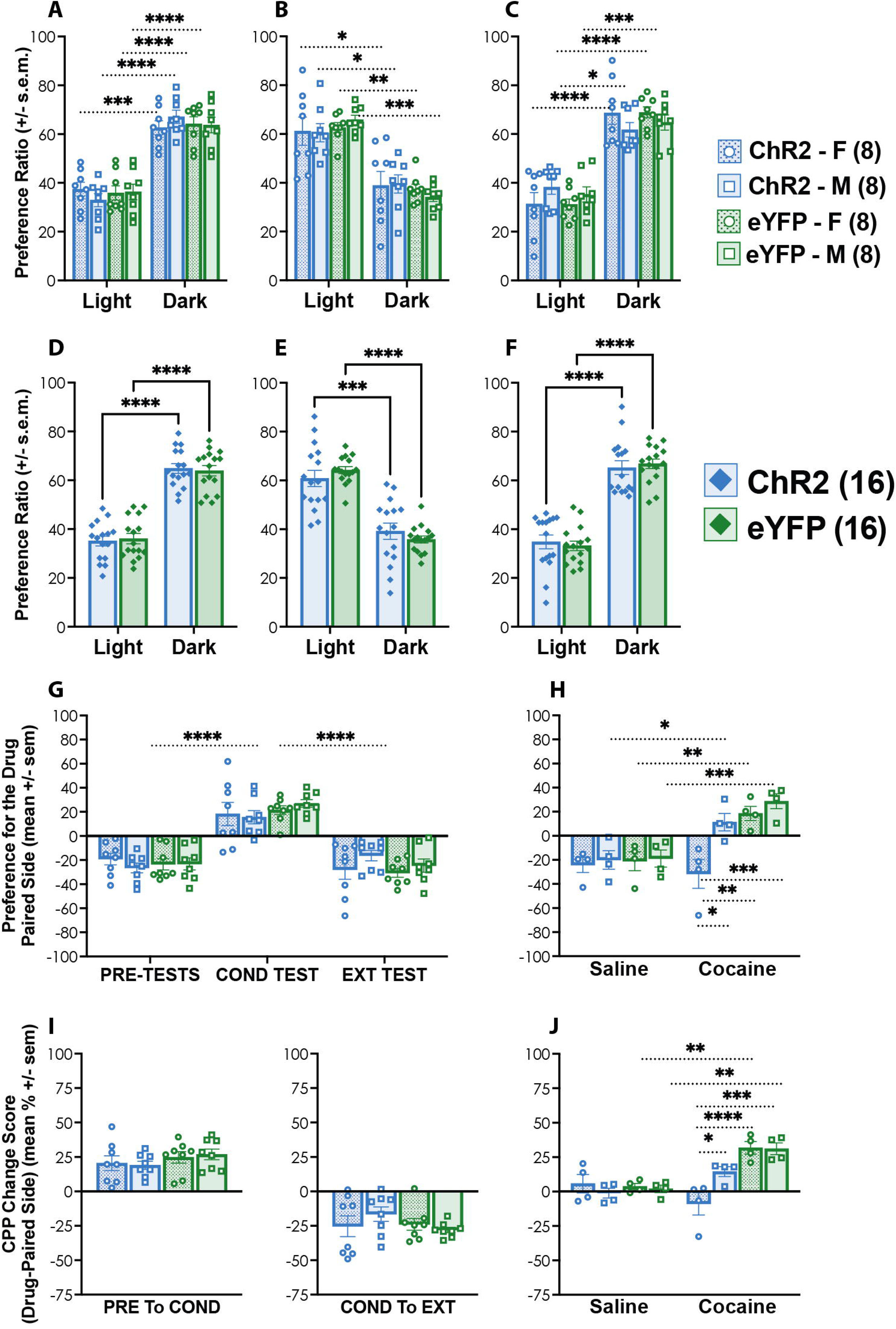
Reactivation of dDG engrams tagged during a first saline exposure attenuates cocaine-primed reinstatement selectively in females. (A–F) Preference ratio across pre-test, conditioning, and extinction sessions. Mice initially preferred the dark side, shifted preference to the cocaine-paired light side following conditioning, and returned to preferring the dark side following extinction. (G) Preference scores across testing phases demonstrating acquisition and extinction of CPP. (H) Cocaine priming reinstated CPP in all groups except female ChR2 mice, suggesting that reactivation of saline-associated ensembles interfered with cocaine-induced reinstatement selectively in females. (I) CPP change scores across pre-test, conditioning, and extinction sessions revealed no group differences. (J) During reinstatement, cocaine increased CPP change scores in all groups except female ChR2 mice, indicating that reactivation of the saline contextual experience attenuated cocaine-induced CPP selectively in females. Data are presented as mean ± s.e.m. *p < 0.05, **p < 0.01, ***p < 0.001, ****p < 0.0001.

**Figure 9.**
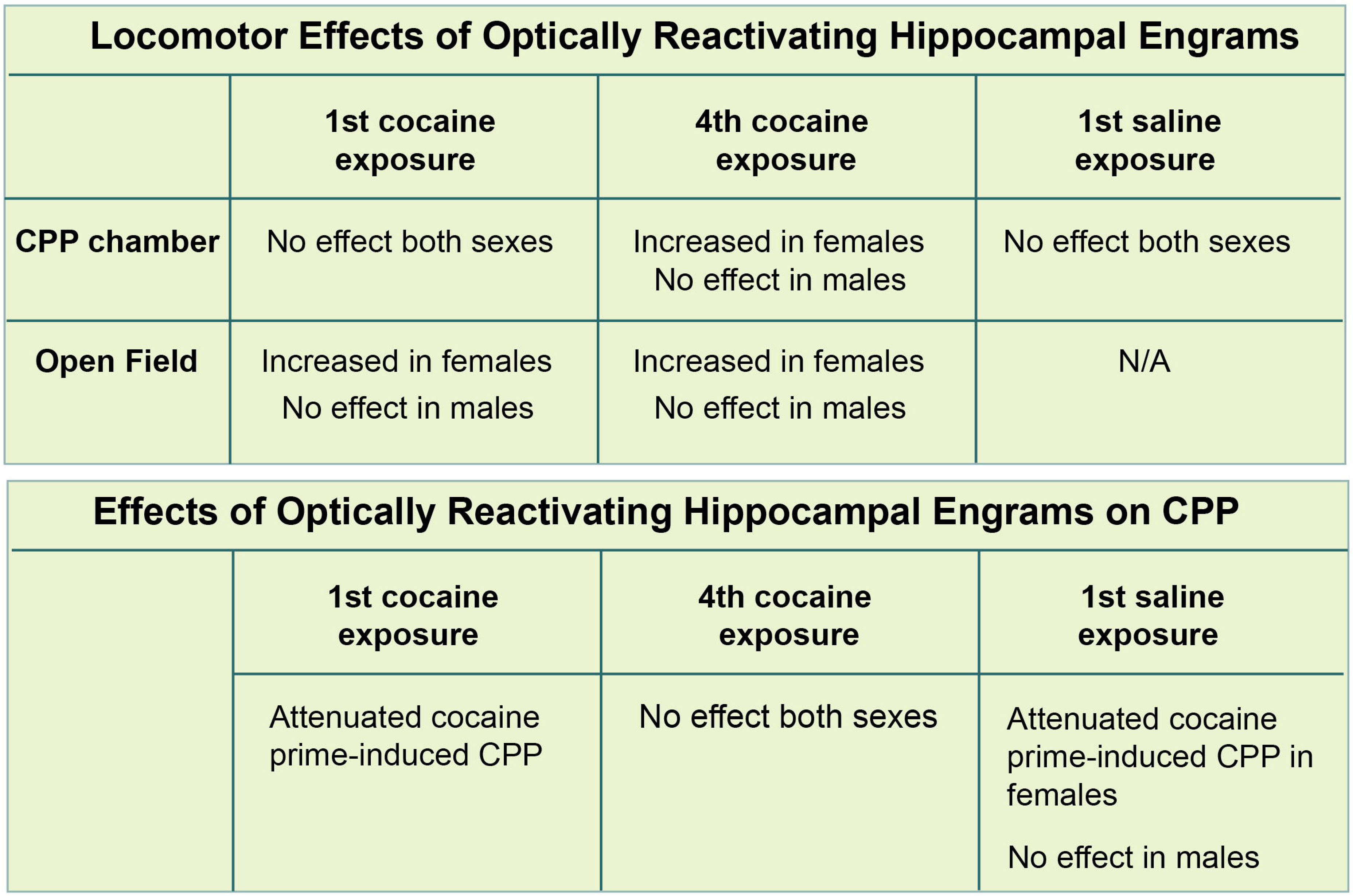
Summary model of how reactivation of distinct dDG engrams influences locomotor and cocaine-seeking behavior. Schematic summary illustrating how reactivation of dDG ensembles tagged during distinct behavioral experiences differentially influences cocaine-seeking. Reactivation of ensembles encoding a first cocaine exposure attenuated cocaine-primed reinstatement, whereas reactivation of ensembles encoding a fourth cocaine exposure did not. Reactivation of saline-associated ensembles also attenuated cocaine-induced CPP selectively in females. Females additionally showed greater sensitivity to the locomotor-inducing effects of cocaine memory reactivation. Together, these findings suggest that early contextual representations may interfere with subsequent cocaine-associated memories and modulate relapse-like behavior.

**Figure 8G** further demonstrates how preference changed across testing (ME:TEST, F_2,56_ = 83.98, p<0.0001, 3-way RM ANOVA). During reinstatement, a cocaine prime reinstated drug-seeking behavior **(Figure 8H)** (DRUG x VIRUS INTERACTION, F_1,24_ = 8.335, p=0.0081; DRUG x SEX INTERACTION, F_1,24_ = 4.533, p=0.0437; 3-way ANOVA) in all groups except the ChR2 female group, which was significantly different than male counterparts (p=0.0181) and both eYFP groups (males: p=0.0003; females: p=0.0034).

There were no group differences in change in preference for the drug-paired side across testing **(Figures 8I)** (*ns*, 2-way ANOVA). During reinstatement **(Figure 8J)**, cocaine increased CPP relative to saline in all groups except the ChR2 females DRUG x VIRUS x SEX INTERACTION, F_1,24_ = 4.528, p=0.0438; 3-way ANOVA) suggesting that reactivation of the saline contextual experience attenuated cocaine-induced CPP selectively in female mice.

Finally, across experiments, for each condition separately, we ran analyses to assess whether locomotor activity (distance and speed) exhibited during the reinstatement test was correlated with preference for the drug paired side, and found that even in the cocaine-eYFP groups it was not (*ns*, Pearson’s correlation) demonstrating a dissociation between locomotor-inducing effects of memory reactivation and CPP.

## 4. DISCUSSION

Together, these findings suggest that reactivation of hippocampal ensembles formed during cocaine experience can modulate subsequent cocaine-seeking behavior. Rather than promoting reinstatement, reactivation of the ensemble tagged during the first cocaine exposure attenuated cocaine-primed relapse-like responding, consistent with the possibility that reactivation engaged competing contextual or affective representations that disrupted expression of the subsequently established drug-associated memory. In contrast, attenuation was not observed following reactivation of the ensemble tagged during the fourth cocaine exposure. This pattern raises the possibility that the influence of cocaine-associated dDG ensembles on later behavior may vary depending on when they are recruited across repeated cocaine experiences, consistent with the idea that early drug-related contextual representations may retain properties that influence subsequent drug seeking (Cutler et al., 2025).

However, because the first- and fourth-exposure conditions were examined in separate experiments and independent cohorts, these findings do not provide a direct comparison of the two populations. Additionally, stronger initial conditioning in the fourth-exposure ChR2 cohort may have influenced subsequent reinstatement. Direct comparison within a common experimental design will be needed to determine whether ensembles recruited at different stages of cocaine experience differ in their capacity to modulate later cocaine seeking.

These findings parallel our previous work (Grella et al., 2022) demonstrating that artificial reactivation of hippocampal representations during fear memory reconsolidation can modify subsequent emotional behavior. In that study, reactivation of dDG representations during fear memory-updating reduced later fear expression, supporting the idea that hippocampal ensemble activation can bias memory expression through interference or updating mechanisms rather than simply driving recall. The present findings extend this framework to cocaine-associated memories, suggesting that reactivation of contextual hippocampal representations may similarly alter the behavioral expression of drug-associated memories during reinstatement testing (Cutler et al., 2025). They also complement previous work demonstrating a causal role for discrete neuronal populations in cocaine-associated memory. Hsiang et al., (2014) for example, showed that a sparse population of neurons within the lateral amygdala was preferentially allocated to a cocaine-cue memory and that manipulating these allocated neurons altered subsequent cocaine CPP expression. Whereas this work established a role for allocated lateral amygdala neurons in cocaine-memory expression, the present study targets experience-dependent dDG ensembles and asks how their artificial reactivation influences cocaine-primed reinstatement following extinction. Our findings therefore extend ensemble-based approaches to the hippocampal contextual component of cocaine-associated memory and suggest that reactivation of these representations can modulate, rather than simply reproduce, subsequent cocaine-seeking behavior.

An important consideration is that activity-dependent TetTag labeling is not restricted exclusively to neurons activated during the cocaine-conditioning episode. Although DOX was removed 42 h before the targeted experience, this interval includes the time required for DOX levels to decline sufficiently to permit tTA-dependent transcription and therefore should not be interpreted as a continuous 42-h labeling period (Ramirez et al., 2013). Moreover, basal neuronal recruitment within the dDG is relatively sparse (Schmidt et al., 2012). Using the same activity-dependent tagging approach, we previously observed labeling of approximately 6–12% of dDG neurons following defined behavioral experiences compared with approximately 2% under home-cage conditions (Grella et al., 2022; Ramirez et al., 2013). Nevertheless, neurons recruited by other components of the experimental experience, including handling, injection, transport, and context exposure, may contribute to the tagged population. For this reason, we refer to these cells as cocaine-associated dDG ensembles rather than assuming that all tagged neurons specifically encode the cocaine-context association.

In addition, because the activity-dependent TetTag approach used here was not restricted to a genetically defined neuronal subtype, we cannot exclude recruitment of local interneurons in addition to DG granule cells. Thus, the present experiments do not establish the precise cellular composition of the tagged population, and the behavioral effects of optical stimulation should be interpreted as resulting from manipulation of an activity-defined dDG ensemble rather than a selectively targeted neuronal subtype. Although c-Fos immunohistochemistry demonstrated increased recruitment of tagged dDG neurons following optical stimulation in ChR2-expressing mice relative to eYFP-only controls, this approach does not establish the fidelity of action potential generation in response to individual light pulses. Direct electrophysiological recordings would be required to assess the temporal precision of optogenetically evoked neuronal activity.

The saline-tagging experiment further tested whether attenuation of cocaine-primed reinstatement required reactivation of an ensemble associated with cocaine exposure, or whether reactivation of an earlier, non-cocaine experience representation could similarly influence subsequent cocaine seeking. Importantly, the saline-tagged ensemble should not be considered a neutral or nonspecific population, as it represents an experience encompassing the experimental context, handling, and injection procedures, but in the absence of cocaine. Unexpectedly, reactivation of this ensemble also attenuated cocaine-primed reinstatement, particularly in females indicating that the behavioral effect of dDG ensemble reactivation does not appear to require cocaine-specific content. Saline-tagged mice exhibited greater ensemble size than cocaine-tagged mice, suggesting that the extent or composition of recruited hippocampal populations may contribute to the observed behavioral effects. This interpretation is consistent with our previous findings (Grella et al., 2022), in which artificial reactivation of larger, non-specific hippocampal cell populations produced stronger behavioral modulation than reactivation of smaller ensembles, suggesting that the magnitude of recruited hippocampal activity itself may influence memory expression. In that study (Grella et al., 2022), we also observed that reactivation of ensembles recruited earlier in an experience produced greater modulation of subsequent fear-memory expression (Grella et al., 2022).

Thus, one possible interpretation is that reactivation of an earlier experience representation competes with the subsequently established cocaine-context representation, thereby interfering with its retrieval or expression during cocaine-primed reinstatement. More broadly, neuronal allocation mechanisms provide a means through which representations of different experiences can interact. Lau et al., (2020) demonstrated that experience-dependent changes in neuronal excitability can promote co-allocation of neurons across temporally proximal events, thereby linking their representations and influencing subsequent memory expression. Although the present experiments do not test co-allocation or memory linking directly, this work supports the broader idea that the neuronal representation of an earlier experience can influence how subsequent experience is represented and behaviorally expressed. In the present study, we hypothesize that artificial reactivation of an earlier dDG representation may similarly influence expression of a subsequently established cocaine-context memory, potentially by biasing retrieval toward the earlier representation or creating competition between the two representations.

Both the saline and first cocaine exposures occurred early in the animals’ experience, when the experimental context and associated procedures were relatively novel. Novelty and uncertainty strongly engage hippocampal and neuromodulatory systems involved in salience encoding and memory allocation (12,23–27) potentially giving these early representations properties that allow their later reactivation to influence expression of subsequently acquired information. Importantly, the ensemble recruited during the first cocaine exposure was tagged before a stable cocaine-context preference had been established and therefore should not be interpreted as representing the fully formed cocaine-context memory subsequently expressed during CPP. Rather, this population may encode multiple features of the initial cocaine experience, including contextual, interoceptive, rewarding, and potentially aversive components. One possibility is therefore that artificial reactivation of this earlier representation introduces information that competes with the subsequently established cocaine-context association during reinstatement. A purely aversive or stress-based account is unlikely to fully explain the present findings, as stress and negative affective states are generally associated with enhanced cocaine-seeking and reinstatement (Erb et al., 1996, 2001; Grella et al., 2014; Mantsch et al., 2016), whereas reactivation of both early saline and early cocaine-associated ensembles attenuated cocaine-primed CPP. Instead, reactivation of earlier contextual or internal-state hippocampal representations compete with, destabilize, or otherwise interfere with expression of the subsequently established cocaine-context association. The particularly robust effect of saline-ensemble reactivation in females further suggests that the behavioral influence of these representations may be sex-dependent and requires further investigation. In contrast, ensembles formed during repeated cocaine exposure may reflect more consolidated or habitual drug-associated representations that are less effective at disrupting reinstatement when reactivated.

The present findings also highlight the importance of contextual representations in cocaine sensitization and reinstatement. Psychostimulant sensitization is highly context dependent (Mattson et al., 2008), and artificial reactivation of hippocampal ensembles likely reinstates components of the contextual state associated with encoding rather than reproducing the pharmacological effects of cocaine itself (Grella et al., 2022). Consistent with this interpretation, optical stimulation alone did not reliably induce CPP reinstatement, despite modulating subsequent cocaine-primed responding. Future studies directly testing reactivation effects across distinct contexts, including contexts outside of the conditioning context, will be important for clarifying the contextual specificity of these effects.

Females were generally more sensitive to the locomotor-activating effects of cocaine memory reactivation and, in some conditions, exhibited stronger attenuation of cocaine-primed CPP (Cailhol & Mormède, 1999; Catalfio et al., 2023; Peart et al., 2022). However, locomotor and CPP effects were dissociable, suggesting that hippocampal ensemble reactivation influences motivational and psychomotor processes through partially distinct mechanisms. Because females were not examined in our previous work (Grella et al., 2022), these findings further emphasize the importance of including sex as a biological variable in studies of memory-based modulation of drug-seeking behavior (Bale & Epperson, 2017; Hunsberger et al., 2025; Shansky & Murphy, 2021).

Importantly, hippocampal ensemble activity was associated with reinstatement-related behavior, supporting a relationship between contextual memory representations and relapse-like responding. Together, these findings suggest that reactivation of hippocampal contextual memories can modulate cocaine-seeking behavior depending on the nature and timing of the encoded experience. Specifically, reactivation of ensembles formed during early experiences appeared to interfere with cocaine-primed reinstatement, whereas ensembles formed following repeated cocaine exposure produced weaker behavioral modulation. A summary of these proposed effects is provided in **Figure 8**.

The present experiments examined the behavioral consequences of artificially reactivating cocaine-associated dDG ensembles but did not test whether these populations are necessary for expression of cocaine CPP. Addressing necessity would require selective inhibition of the tagged population during memory retrieval. Notably, artificial activation of the tagged ensembles did not itself produce CPP or substitute for a cocaine prime; rather, reactivation modulated cocaine-primed reinstatement, with activation of earlier-recruited ensembles interfering with expression of the previously established cocaine-context preference. Thus, the present findings support a role for these dDG representations in modulating subsequent cocaine-seeking behavior rather than establishing their necessity for cocaine-memory retrieval.

## Ethics Approval

All experimental procedures were approved by the Institutional Animal Care and Use Committee (IACUC) at Loyola University Chicago and were conducted in accordance with the National Institutes of Health Guide for the Care and Use of Laboratory Animals.

## Data Availability Statement

The datasets generated and/or analyzed during the current study are available from the corresponding author on reasonable request.

## Primary funding

This work was funded by startup funds.

## Clinical trial registration details

N/A

## Authors’ Contributions

SLG conceived the study. SLG designed the CPP experiments; SLG LHE, LFP, MRW, and SAA designed the open field experiment. LHE, LFP, MRW, MVB, WFW, MC, SAA, AS, SCdV, and SLG collected the data. SLG analyzed the data. SLG LHE, LFP, MRW, and MVB wrote the first draft. LHE, LFP, MRW, MVB, WFW, MC, SAA, AS, SCdV, and SLG edited the final manuscript. All authors approved the final version of the manuscript.

## Acknowledgements

We would like to thank Dr. Stephan Steidl, Joe Schluep, and Monica Mackey for technical guidance. We thank Beata Czesny and the animal care staff for their support. We also thank the NIH NIDA Drug Supply Program for providing cocaine hydrochloride.

## List of Abbreviations

AAV: Adeno-associated virus
ANOVA: analysis of variance
AP: anterior / posterior
ChR2: Channelrhodopsin-2
COND: Conditioning
CPP: Conditioned Place Preference
DAPI: 4′,6-diamidino-2-phenylindole
dDG: Dorsal Dentate Gyrus
DOX: Doxycycline
DV: Dorsal / ventral
EXT: Extinction
eYFP: Enhanced yellow fluorescent protein
ME: Main effect
ML: Medial / lateral
NIDA: National Institute on Drug Addiction
NIH: National Institute of Health
PBS: Phosphate buffered saline
PBST: Phosphate buffered saline with 2% Triton
PFA: Paraformaldehyde
PRE: Pre-tests
TRE: Tetracycline response element
tTA: Tetracycline transactivator protein

